# PDLIM2 Repression: A Common Mechanism in Viral Lung Infection

**DOI:** 10.1101/2025.09.12.675949

**Authors:** Feng Gao, Yongzheng W. Chen, Steven D. Shapiro, Gutian Xiao, Zhaoxia Qu

## Abstract

**Background:** PDLIM2, a PDZ-LIM domain-containing protein expressed highest in the lung and immune cells, serves as a unique tumor suppressor and immune modulator, mainly by turning off the activation of the master transcription factors NF-κB and STAT3. While its role in cancer is established, the involvement of PDLIM2 in viral infection remains unclear.

**Results:** Here, we analyzed public gene expression data of blood leukocytes, bronchoalveolar lavage cells, and lung tissues from uninfected healthy humans and those infected with the respiratory virus SARS-CoV-2 or influenza. We found that PDLIM2 expression was repressed by viral infection, and notably, this repression correlated with the severity of infectious diseases. Consistently, the expression level of PDLIM2 was negatively associated with NF-κB and STAT3 activity across a diverse range of cell types, such as macrophages, monocytes, neutrophils, T cells, alveolar type 1 and 2 epithelial cells, airway epithelial cells, and fibroblasts. Accordingly, cells with low PDLIM2 expression exhibited aberrant activation of signaling pathways essential for cellular functions and immune responses.

**Conclusions:** These findings highlight PDLIM2 repression as a common mechanism underlying human viral infectious diseases and suggest PDLIM2 as a potential biomarker and therapeutic target for disease prognosis, prevention, and treatment.

## Background

Lung infection is a leading cause of hospitalization and death in the United States and worldwide [1–3]. And it causes significantly higher morbidity and mortality in individuals with pre-existing conditions, such as lung cancer, chronic obstructive pulmonary disease (COPD), and interstitial lung disease (ILD)/pulmonary fibrosis (PF) [4–6]. In line with the high risk and high death rate of lung infection, unfortunately, our understanding of lung infectious disease, and especially its molecular link with lung cancer and other lung diseases, remains far from complete.

Recent studies identify the PDZ-LIM domain-containing protein PDLIM2 as a key molecular checkpoint in preventing lung cancer, COPD, and ILD/PF [7–11]. PDLIM2, also known as mystique or SLIM, is ubiquitously expressed under physiological conditions, with the highest level in the lung [12–14]. At the cellular level, it is expressed abundantly in various epithelial and immune cells, particularly lung epithelial cells and myeloid cells [7–11, 14, 15]. However, its expression is repressed in the lung of patients with lung cancer, COPD and/or ILD/PF, and this repression is associated with disease severity and poor prognosis and survival [7–10]. In mice, global deletion or selective obliteration of PDLIM2 from lung epithelial cells or myeloid cells renders animals highly susceptible to spontaneous and induced lung tumors as well as to lung injury and death by the endotoxin lipopolysaccharide (LPS) [7, 8, 11]. Notably, PDLIM2 restoration by nano-delivery of exogenous PDLIM2 (nanoPDLIM2) or pharmacological induction of endogenous PDLIM2 shows promising therapeutic activities and improves the efficacy of chemotherapy and immunotherapy in the mouse models of lung cancer [7, 8, 10, 15].

PDLIM2 exerts its tumor suppressor role and immune regulation effect through multiple mechanisms [16–19]. The major one is to terminate the activation of nuclear factor kappaB (NF-κB) and signal transducer and activator of transcription 3 (STAT3) by promoting ubiquitination and proteasomal degradation of activated RelA (the prototypical NF-κB member, also known as p65) and STAT3 in the nucleus [7–10, 20–26]. NF-κB and STAT3 are master transcription factors that control inflammation, metabolic reprogramming, and cell survival; and their persistent activation plays causative roles in the pathogenesis of various human diseases, including lung cancer, COPD, ILD/PF, and infections [7–10, 27–39]. More recent mechanistic studies show that instead of being claimed before as a ubiquitin ligase (E3), PDLIM2 acts as a novel ubiquitin ligase enhancer (E5) to stabilize and chaperone the E3 SCF^β-TrCP^ to nuclear RelA, ubiquitinating it for proteasomal degradation [14, 26, 40, 41].

In this manuscript, we examined the possible involvement and clinical relevance of PDLIM2 in lung viral infection. We exploited public gene microarray and next-generation sequencing (NGS) data of peripheral blood leukocytes, bronchoalveolar lavage (BAL) cells, and lung tissues from uninfected ‘healthy’ humans and those infected with the respiratory virus severe acute respiratory syndrome coronavirus 2 (SARS-CoV-2) or influenza virus (IV). SARS-CoV-2 and IV are respectively the etiological agents of coronavirus disease 2019 (COVID-19) and influenza (commonly known as the flu), two of the most common respiratory illnesses that are highly contagious and often cause severe lung injury and death, particularly among vulnerable populations [1–3].

## Methods

### Data source and selection

All data were from the public Gene Expression Omnibus (GEO) repository that contains various microarray, bulk RNA-sequencing (RNA-seq), and single-cell RNA-seq (scRNA-seq) data. A search for multiomics data from human patients with viral lung infections yielded a list of datasets on IV and SARS-CoV-2 infections. The generated datasets were subjected to routine quality control analysis and PDLIM2 expression examination. Those with good intrinsic quality and comprehensive representation of cell responses were selected, and the samples/cells that were PDLIM2-positive were then processed with the analysis pipelines described below. GEO datasets for the final analyses are listed in Supplementary Table 1. The publications originally associated with these datasets were also listed [42–48].

### Gene microarray and bulk RNA-seq data analysis pipeline

These data were analyzed in RStudio in the way GEO2R performs. GEO2R, an online tool on the NCBI GEO website, offers data that has undergone alignment, quantification, and normalization processes by the original submitters or by NCBI’s own pipelines, and allows for differential gene expression analysis of microarray and, in a beta version, bulk RNA-seq data. In brief, after a GEO Series was imported into RStudio, experimental groups (e.g., disease vs. control) were defined, and samples were assigned to these groups. Gene expression data of PDLIM2 and other genes were fetched along with their metadata. Statistical packages like Limma for microarray data and DESeq2 for RNA-seq (using NCBI-computed raw counts) were used to identify genes significantly altered between the defined groups. Results were then visualized with plots and heatmaps. The R scripts for the analysis were stored on GitHub (https://github.com/Xiao-Qu-Lab/PDLIM2-in-lung-viral-infection/blob/main/GSE101702.R; and https://github.com/Xiao-Qu-Lab/PDLIM2-in-lung-viral-infection/blob/main/GSE157103.R).

### Single-cell RNA-seq data analysis pipeline

These data were analyzed in Seurat, a popular R package for scRNA-seq data analysis. As scRNA-seq data in NCBI repository were deposited in variable formats and stages, our analysis always started with the processed data if available, and if not, a new analysis was carried out to recapitulate the paper’s results with the settings as defined in the paper’s methods. A full typical workflow involves: 1) Importing gene expression matrix and metadata to Seurat and quality control (QC) to filter low-quality cells. 2) Normalization and scaling to account for technical variations. 3) Dimensionality reduction (e.g., PCA, UMAP/t-SNE) to visualize cell relationships in lower dimensions. 4) Clustering to group cells with similar gene expression profiles, identifying distinct cell populations. 5) Marker gene identification to find genes uniquely expressed in each cluster, aiding cell type annotation. 6) Cell assignments to disease and control groups, with the former further divided into 2 disease subgroups based on the levels of the expression of PDLIM2. 7) Performing differential gene expression with the Wilcoxon Rank Sum test among the groups defined above. 8) Exporting gene expression data of PDLIM2 and other gene groups and differentially expressed genes (DEGs) to the local disk. 9) Gene expression data were visualized with plots and heatmaps, and DEGs were imported to Qiagen ingenuity pathway analysis (IPA) for pathway and functional analyses. The R scripts for the analysis were stored on GitHub (https://github.com/ Xiao-Qu-Lab/PDLIM2-in-lung-viral-infection/blob/main/GSE243629.R; https://github.com/ Xiao-Qu-Lab/PDLIM2-in-lung-viral-infection/blob/main/GSE145926.R; and https://github.com/ Xiao-Qu-Lab/PDLIM2-in-lung-viral-infection/blob/main/GSE171524.R).

### Pathway activation score

The component and target genes of a specific signaling pathway were collected from Qiagen and Reactome, and other online resources, and their expressions were summed up and used as the activation score for the pathway. The source and the list of the genes for the pathways studied can be found in the R scripts stored on the GitHub above.

### PDLIM2 level-dependent pathway and function analysis

Besides the direct comparison between the disease and the control, samples from the disease were subset into PDLIM2-low and -high groups using the median as the divider, and differential gene expression was performed from their respective comparison to the control. The DEGs derived were imported into Qiagen IPA for pathway and functional analyses. Comparison analysis was done between PDLIM2 low and high conditions, and the differential activation of the pathway with significant P-values from both conditions was calculated as z-score deviation between them.

### Statistics

Student’s t-test (2-tailed, unpaired) and ordinary 1-way ANOVA were used to assess the significance of differences between two groups and among groups of more than two, respectively [49–51]. Bars on a box plot represent the minimum, lower quartile, median, upper quartile, and maximum, respectively. The Correlation Coefficient (R) for the trendline was calculated by dividing the covariance of the x and y axis variables by the product of their standard deviations. The P-value in IPA pathway analysis was calculated using Fisher’s Exact Test. The z-score in IPA pathway analysis was calculated by comparing the observed gene expression changes in a dataset to the expected directional changes stored in the knowledge base, providing a statistical measure for the predicted activation or inhibition of a pathway. P-values less than 0.05 were considered statistically.

### Data and code access

All R scripts used for analysis were stored on GitHub (https://github.com/Xiao-Qu-Lab/PDLIM2-in-lung-viral-infection) and are freely accessible to the public.

## Results

### PDLIM2 repression by lung viral infection and its association with disease severity

We first analyzed human peripheral blood leukocyte gene microarray data on influenza and bulk RNA-seq data on COVID-19, because they may be used as a noninvasive and easy-to-perform assay for the prognosis of those infectious diseases. As expected, PDLIM2 was easily detected in the blood leukocytes from uninfected healthy people by either microarray (Fig. 1A) or bulk RNA-seq (Fig. 1B). However, its expression was repressed in patients with influenza or COVID-19. Notably, the extent of PDLIM2 repression was associated with increased severity in both diseases.

**Fig. 1:**
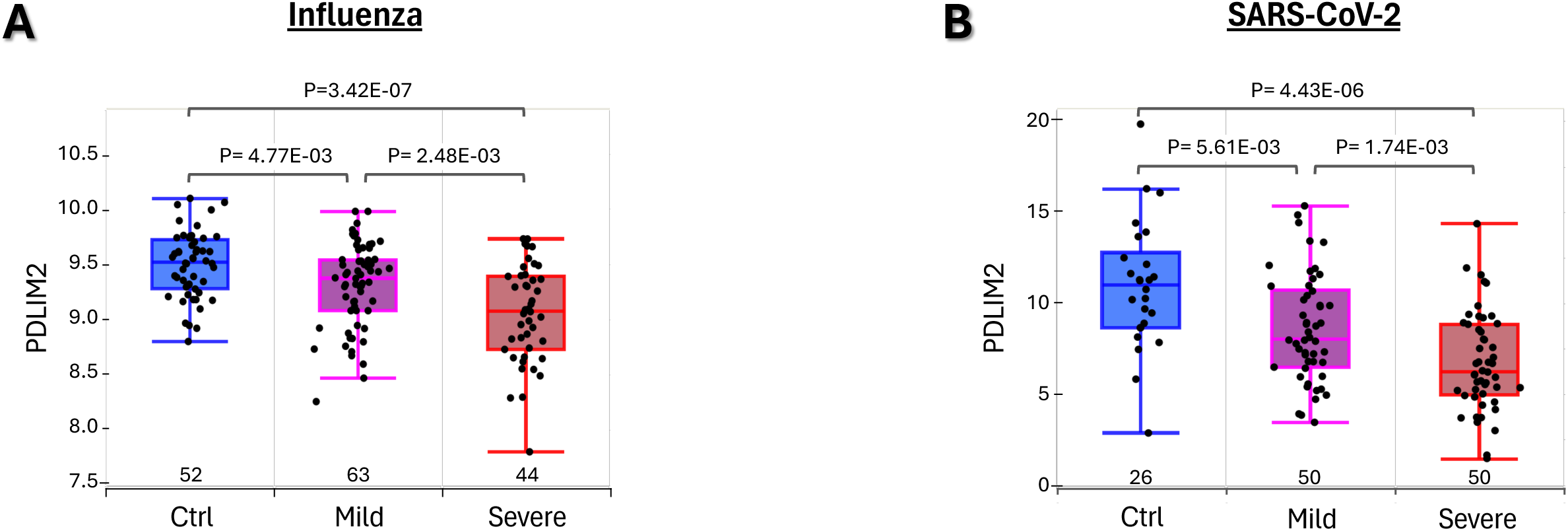
PDLIM2 repression by lung viral infection and its association with disease severity. (A) Gene microarray data analysis showing a negative association of PDLIM2 repression in human peripheral blood leukocytes with influenza severity. (B) Bulk RNA-seq data analysis showing a negative association of PDLIM2 repression in human peripheral blood leukocytes with COVID-19 severity. Sample numbers (N) are listed above the x-axis. Each data point represents one patient. Data is presented as box plots. P-values are obtained from ANOVA or Student’s *t* test.

### Association of PDLIM2 repression with pathological activation and death of neutrophils

Following the promising results, we analyzed scRNA-seq data to examine PDLIM2 expression in peripheral neutrophils, as neutrophils are the most abundant leukocytes in human blood and serve as the first line of defense against infection. In response to lung infection, they move quickly and exponentially to the lung to kill invading pathogens and drive inflammation via several major related mechanisms, including phagocytosis, degranulation, oxidative burst, and release of neutrophil extracellular traps (NETs), a process called NETosis. They also secrete pro-inflammatory cytokines and chemokines, promoting inflammation. After pathogen clearance, neutrophils produce anti-inflammatory cytokines and undergo non-inflammatory apoptosis to be removed mainly by macrophages, leading to the resolution of inflammation and the restoration of tissue homeostasis. Dysregulation of those functions of neutrophils damages lung tissues and promotes disease severity (Supplementary Table 2).

Indeed, PDLIM2 expression was significantly lower in neutrophils from influenza patients in comparison to those from uninfected healthy people (Fig. 2A). Consistent with its known functions, PDLIM2’s expression levels were negatively associated with the activation scores of NF-κB and STAT3 in neutrophils (Fig. 2B). Accordingly, numerous target genes of NF-κB and STAT3 were predominantly more activated in patients’ cells (Fig. 2C). Besides NF-κB and STAT3, many other signaling pathways were detrimentally activated in patients’ neutrophils (Supplementary Fig. 1). Some of them were deregulated also in a manner dependent on PDLIM2 repression, as evidenced by their different activations in the PDLIM2-low versus -high cells of patients and by the association between the activities of these signaling pathways and PDLIM2 expression level (Fig. 4D-4F). Notably, several pathways propelling inflammatory responses, including neutrophil degranulation, necroptosis, NETosis, cellular stress, senescence, and cyclophilin signaling pathways, were increased, whereas those resolving inflammation, such as interleukin-10 (IL-10) signaling were decreased in PDLIM2-low cells. The antiviral interferon (IFN) signaling pathway and the master CXCR4 bone marrow retention signaling of neutrophils were also decreased in PDLIM2-low cells.

**Fig. 2:**
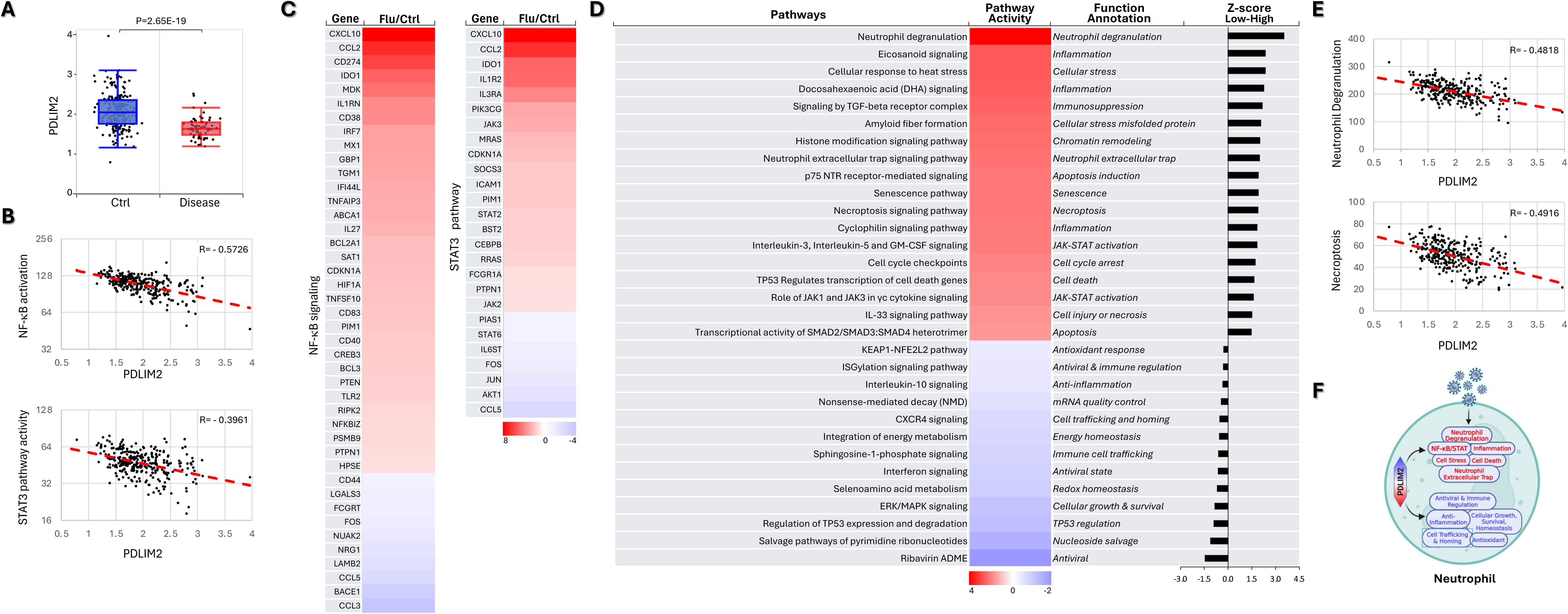
Association of PDLIM2 repression with pathological activation and death of neutrophils. (A) scRNA-seq data analysis showing PDLIM2 repression in the blood neutrophils of influenza patients. Data is presented as box plots. P-values are obtained from Student’s *t* test. (B) Gene and pathway correlation showing negative associations of PDLIM2 expression with activations of the NF-κB and STAT3 signaling pathways in human blood neutrophils. (C) Heatmap showing differential expression of NF-κB and STAT3 target genes in the blood neutrophils of influenza patients in comparison to uninfected ‘healthy’ controls. (D) Heatmap showing differential activation of signaling pathways in PDLIM2-low vs. -high expressing blood neutrophils in influenza patients. (E) Gene and pathway correlation showing negative associations of PDLIM2 expression with the activation of the neutrophil degranulation and necroptosis signaling pathways in human blood neutrophils. (F) Cartoon summary listing the main signaling pathways differentially activated in PDLIM2-low vs. -high expressing blood neutrophils in influenza patients.

### Association of PDLIM2 repression with hyper-activation of monocytes

Upon lung infection, peripheral monocytes, like neutrophils, rapidly migrate to the lung, assisting viral clearance mainly through differentiating into highly inflammatory macrophages and presenting viral antigens to boost adaptive immunity. Monocytes also produce pro-inflammatory cytokines and substances to enhance inflammation. Not surprisingly, overactivation of monocytes leads to excessive inflammation and lung damage, and pathology (Supplementary Table 3).

Same as in neutrophils, PDLIM2 expression was repressed in monocytes from influenza patients compared to those from uninfected healthy humans; and this repression was associated with ultra-activation of NF-κB and STAT3 (Fig. 3A-3C). Numerous other signaling pathways critical for monocytes as the contributors of inflammatory responses were also overwhelmingly activated in patients (Supplementary Fig. 2). Like NF-κB and STAT3, many of these signaling pathways were over-activated in a manner depending on PDLIM2 repression, such as oxidative stress, senescence-associated secretory phenotype (SASP), high mobility group box 1 (HMGB1), triggering receptor expressed on myeloid cells 1 (TREM1), and pathogen-induced cytokine storm signaling pathways (Fig. 3D-3F).

**Fig. 3:**
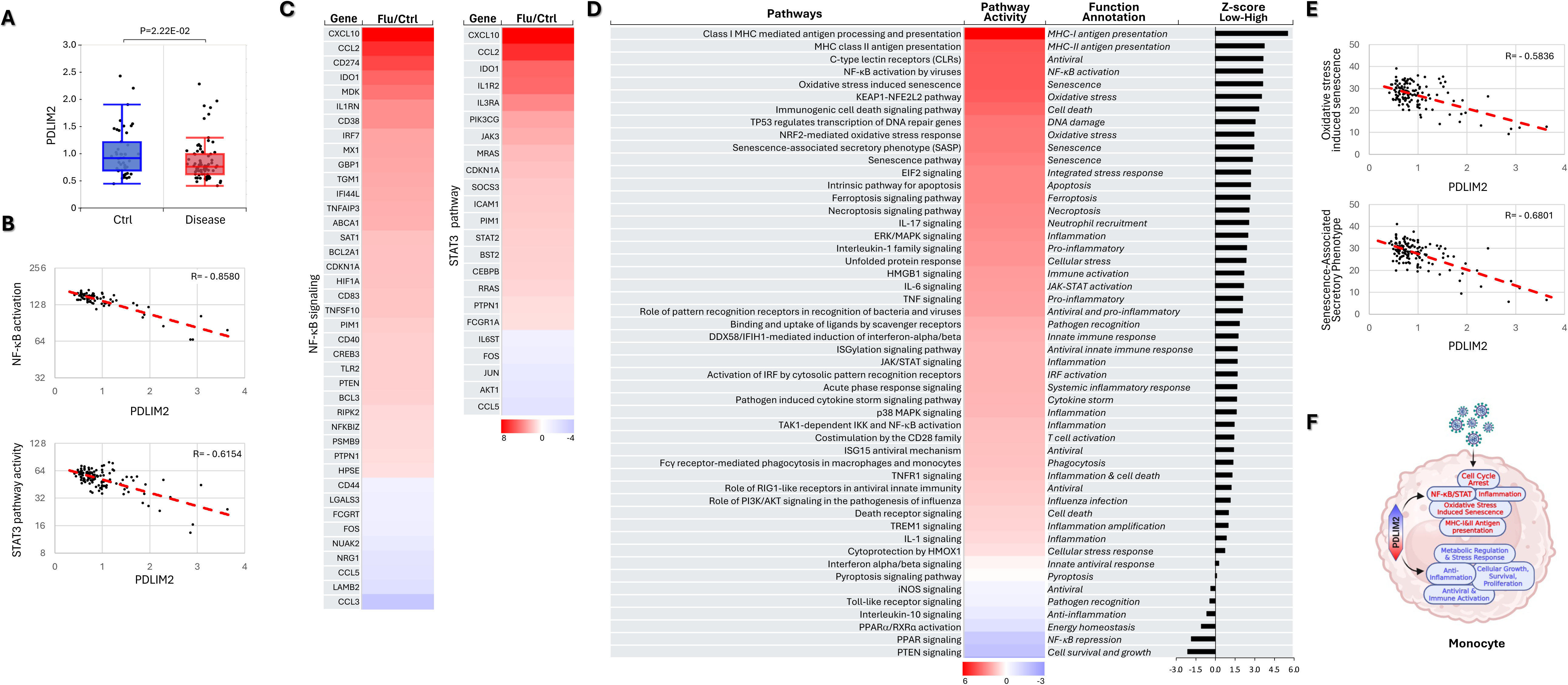
Association of PDLIM2 repression with hyper-activation of monocytes. (A) scRNA-seq data analysis showing PDLIM2 repression in the blood monocytes of influenza patients. P-values are obtained from Student’s *t* test. (B) Gene and pathway correlation showing negative associations of PDLIM2 expression with activations of the NF-κB and STAT3 signaling pathways in human blood monocytes. (C) Heatmap showing differential expression of NF-κB and STAT3 target genes in the blood monocytes of influenza patients in comparison to uninfected ‘healthy’ controls. (D) Heatmap showing differential activation of signaling pathways in PDLIM2-low vs. -high expressing blood monocytes in influenza patients. (E) Gene and pathway correlation showing negative associations of PDLIM2 expression with the activation of the oxidative stress and SASP signaling pathways in human blood monocytes. (F) Cartoon summary listing the main signaling pathways differentially activated in PDLIM2-low vs. -high expressing blood monocytes in influenza.

**Fig. 4:**
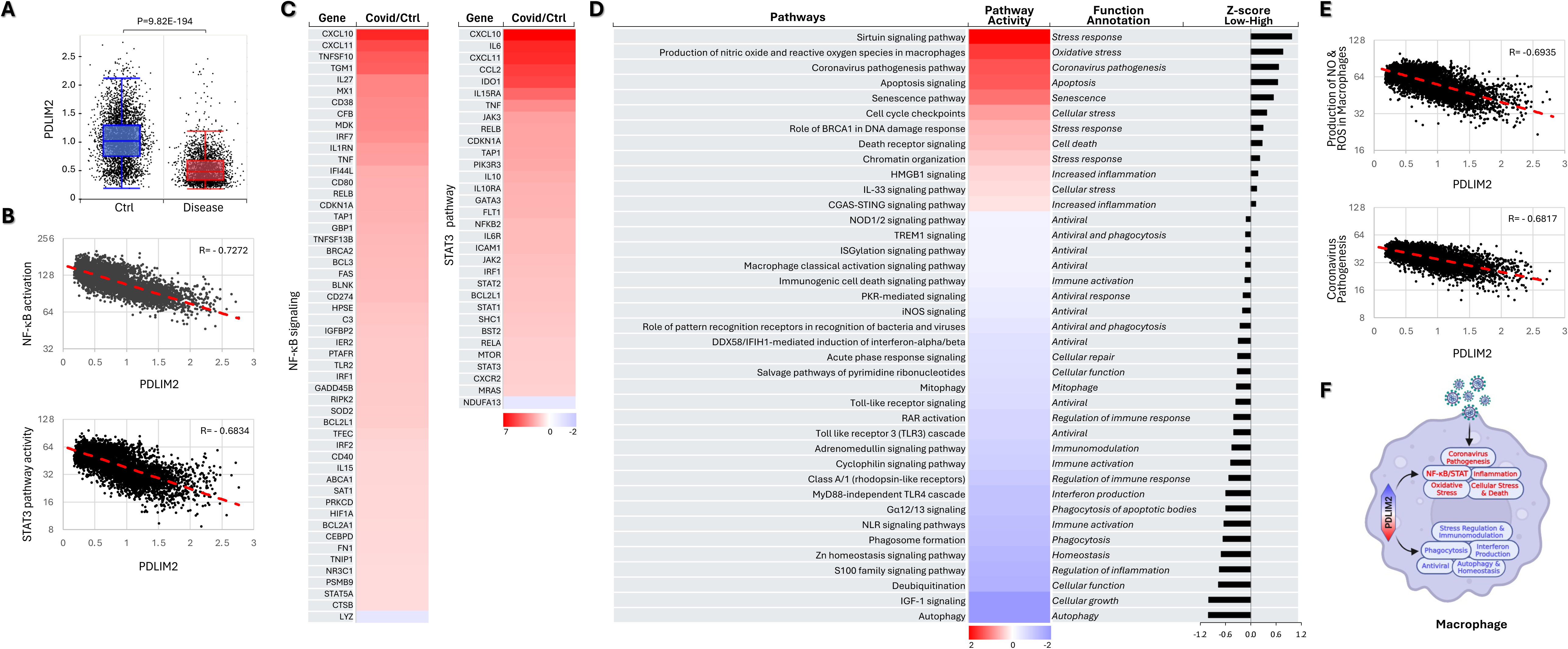
Association of PDLIM2 repression with pathogenic activation of lung macrophages. (A) scRNA-seq data analysis showing PDLIM2 repression in the lung macrophages of COVID-19 patients. P-value was obtained from Student’s *t* test. (B) Gene and pathway correlation showing negative associations of PDLIM2 expression with activations of the NF-κB and STAT3 signaling pathways in human lung macrophages. (C) Heatmap showing differential expression of NF-κB and STAT3 target genes in the lung macrophages of COVID-19 patients in comparison to uninfected controls. (D) Heatmap showing differential activation of signaling pathways in PDLIM2-low vs. -high expressing lung macrophages in COVID patients. (E) Gene and pathway correlation showing negative associations of PDLIM2 expression with the activation of the production of NO & ROS and coronavirus pathogenesis signaling pathway in human lung macrophages. (F) Cartoon summary listing the main signaling pathways differentially activated in PDLIM2-low vs. -high expressing lung macrophages in COVID-19 patients.

### Association of PDLIM2 repression with pathogenic activation of lung macrophages

Macrophages, the primary phagocytes, are the most abundant immune cells within the lung. Particularly, those residing in and patrolling the alveoli serve as the first immune cell responder to lung infection, killing inhaled pathogens and turning on inflammation and resolving it to promote lung tissue healing after pathogen clearance. Dysregulation in this functional switch of those key sentinels leads to uncontrolled inflammation and causes severe lung damage and often death (Supplementary Table 4).

Analysis of the scRNA-seq data of BAL cells showed that many signaling pathways were activated differently in the lung macrophages from COVID-19 patients compared to those from uninfected controls (Supplementary Fig. 3). Importantly, we identified a drastic PDLIM2 repression and a strong negative association of PDLIM2 expression with NF-κB and STAT3 activation in these culprit cells of the fatal pathogenesis (Fig. 4A-4C). In addition to NF-κB and STAT3, several other pro-inflammatory signaling pathways, such as oxidative stress, senescence, HMGB1, and coronavirus pathogenesis pathways, were hyper-activated in a PDLIM2 repression-dependent manner (Fig. 4D-4F). On the contrary, those involved in viral clearance and inflammation resolution, including phagocytosis, autophagy, insulin-like growth factor (IGF), and S100 family signaling, were hypo-activated, which also depended on PDLIM2 repression. Of note, autophagy also serves as an intrinsic checkpoint restricting NF-κB activities [52–55].

### Association of PDLIM2 repression with T-cell exhaustion and senescence

During lung infection and especially when myeloid cells (macrophages and neutrophils) are insufficient to control it, T cells are activated and migrate into the lung where they eliminate infected cells and release cytokines that help control the infection. Similarly, T cells may also contribute to lung inflammation and injury, including those caused by viruses like SARS-CoV-2 (Supplementary Table 5 and Supplementary Fig. 4). Same as myeloid cells, T cells in COVID-19 patients expressed PDLIM2 at a much lower level and had high NF-κB and STAT3 activation (Fig. 4A-4D). On the other hand, Rho GDP-dissociation inhibitor (RhoGDI) signaling, which plays essential roles in various cellular functions, including T cell activation and migration, was lower in T cells with low PDLIM2 expression in comparison to those with high PDLIM2 expression. Remarkably, T cells with low PDLIM2 expression were more exhausted and with an increased SASP.

### Association of PDLIM2 repression with viral endocytic entry and immunogenic cell deaths of alveolar type 2 epithelial cells

Alveolar type 2 (AT2) cells, one of the two major types of epithelial cells lining the pulmonary alveoli, are the primary source of pulmonary surfactants and adult tissue stem/progenitor cells, thereby critical for respiratory homeostasis and lung tissue repair after injury. However, they are also the main targets of lung infection, and their dysfunction and damage may lead to various lung conditions and often death. Despite being non-immune cells, AT2 cells possess unique immunologic properties and act as active participants in both the innate and adaptive immune responses, influencing the severity and outcome of diseases like COVID-19 (Supplementary Table 6).

Analysis of scRNA-seq data of human lung tissues revealed that a plethora of signaling pathways were altered within the AT2 cells of COVID-19 patients (Supplementary Fig. 5). Many of them and particularly those involved in the fundamental activities of AT2 cells were deregulated in association with PDLIM2 repression, such as NF-κB, STAT3, surfactant metabolism, oxidative stress and immune regulation pathways (Fig. 6A-6D). Intriguingly, endocytic viral entry, senescence, and immunological cell death pathways were markedly increased, whereas RhoGDI, peroxisome proliferator-activated receptor (PPAR), and IL-10 signaling pathways were decreased in AT2 cells in which PDLIM2 was repressed (Fig.6D-6F). Note, PPAR signaling, like RhoGDI signaling, is involved in the cell repair, maintenance, and differentiation of AT2 and many other cells. PPAR signaling can also repress NF-κB.

**Fig. 5:**
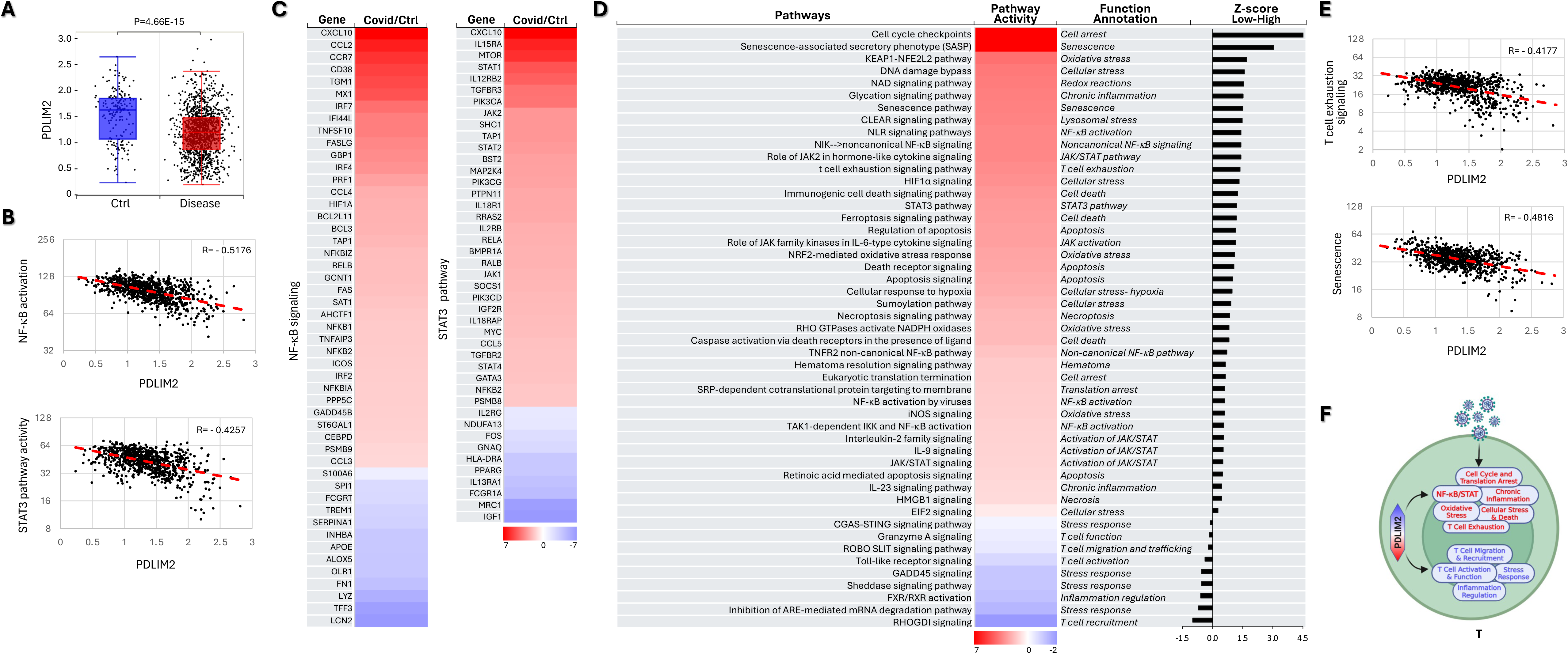
Association of PDLIM2 repression with T-cell exhaustion and senescence. (A) scRNA-seq data analysis showing PDLIM2 repression in the lung T cells of COVID-19 patients. P-value was obtained from Student’s *t* test. (B) Gene and pathway correlation showing negative associations of PDLIM2 expression with activations of the NF-κB and STAT3 signaling pathways in human lung T cells. (C) Heatmap showing differential expression of NF-κB and STAT3 target genes in the lung T cells of COVID-19 patients in comparison to uninfected controls. (D) Heatmap showing differential activation of signaling pathways in PDLIM2-low vs. -high expressing lung T cells in COVID-19 patients. (E) Gene and pathway correlation showing negative associations of PDLIM2 expression with the activation of the T-cell exhaustion and senescence signaling pathways in human lung T cells. (F) Cartoon summary listing the main signaling pathways differentially activated in PDLIM2-low vs. -high expressing lung T cells in COVID-19 patients.

**Fig. 6:**
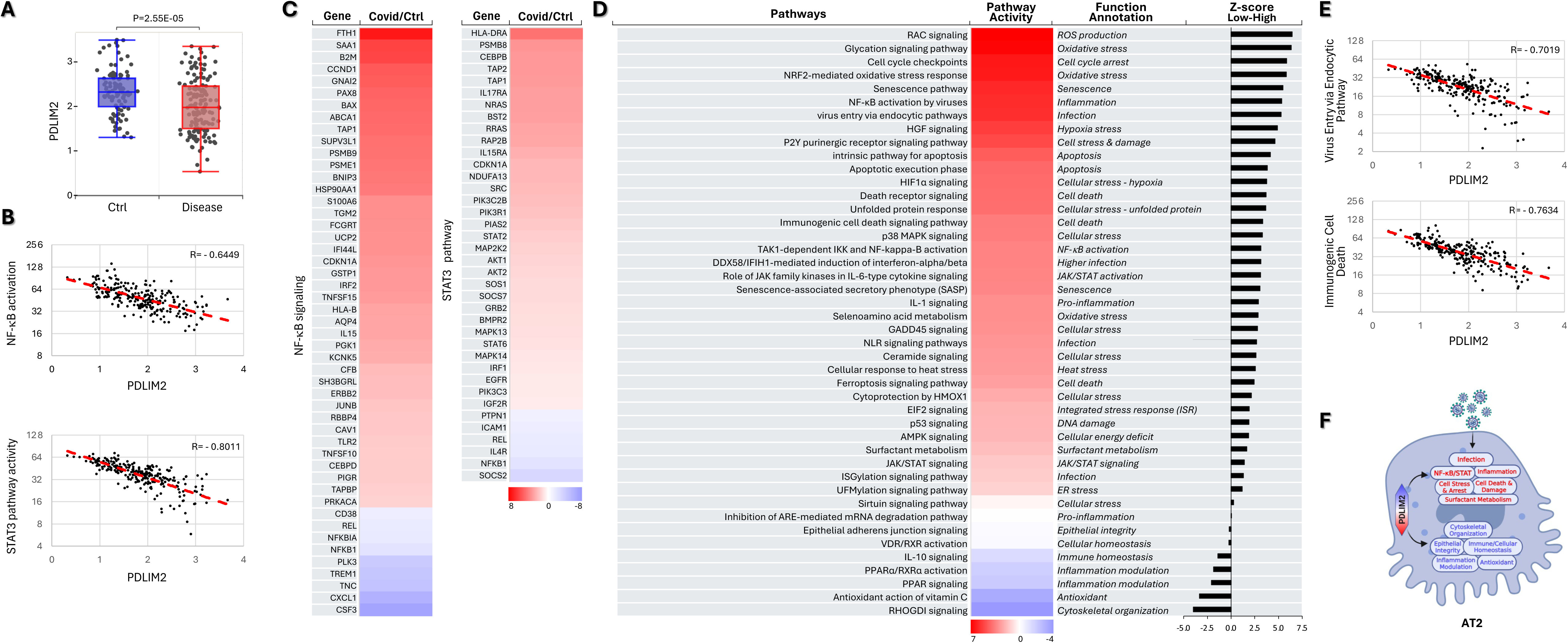
Association of PDLIM2 repression with viral endocytic entry and immunogenic cell deaths of AT2 cells. (A) scRNA-seq data analysis showing PDLIM2 repression in the AT2 cells of COVID-19 patients. P-value was obtained from Student’s *t* test. (B) Gene and pathway correlation showing negative associations of PDLIM2 expression with activations of the NF-κB and STAT3 signaling pathways in human AT2 cells. (C) Heatmap showing differential expression of NF-κB and STAT3 target genes in the AT2 cells of COVID-19 patients in comparison to uninfected ‘healthy’ controls. (D) Heatmap showing differential activation of signaling pathways in PDLIM2-low vs. -high expressing AT2 cells in COVID-19 patients. (E) Gene and pathway correlation showing negative associations of PDLIM2 expression with the activation of the endocytic viral entry and immunological cell death signaling pathways in human AT2 cells. (F) Cartoon summary listing the main signaling pathways differentially activated in PDLIM2-low vs. -high expressing AT2 cells in COVID-19 patients.

### Association of PDLIM2 repression with alveolar type 1 epithelial cell damage

Alveolar type 1 (AT1) cells, the major type of alveolar epithelial cells other than AT2 cells, are responsible for gas exchange in the lung. They also play a significant role in lung immunity, interacting directly with immune cells and releasing immune mediators like cytokines and chemokines. AT1 cell dysfunction or damage is a major factor in various lung conditions, including infectious diseases like COVID-19, and often leads to severe outcomes and death (Supplementary Table 7 and Supplementary Fig. 6).

Like AT2 cells, AT1 cells in COVID-19 patients had a PDLIM2 repression-dependent hyper-activation of NF-κB and STAT3 (Fig. 7A-7D). Other over-activated signaling pathways dependent on PDLIM2 repression included those involved in senescence, cell cycle arrest, oxidative and cellular stress, and immunological cell death (Fig. 7D-7F). Interestingly, RhoGDI signaling pathway and chaperone-mediated autophagy (CMA), another signaling vital for AT1 cell repair, maintenance, and function, were decreased significantly in patients’ AT1 cells with low PDLIM2 expression.

**Fig. 7:**
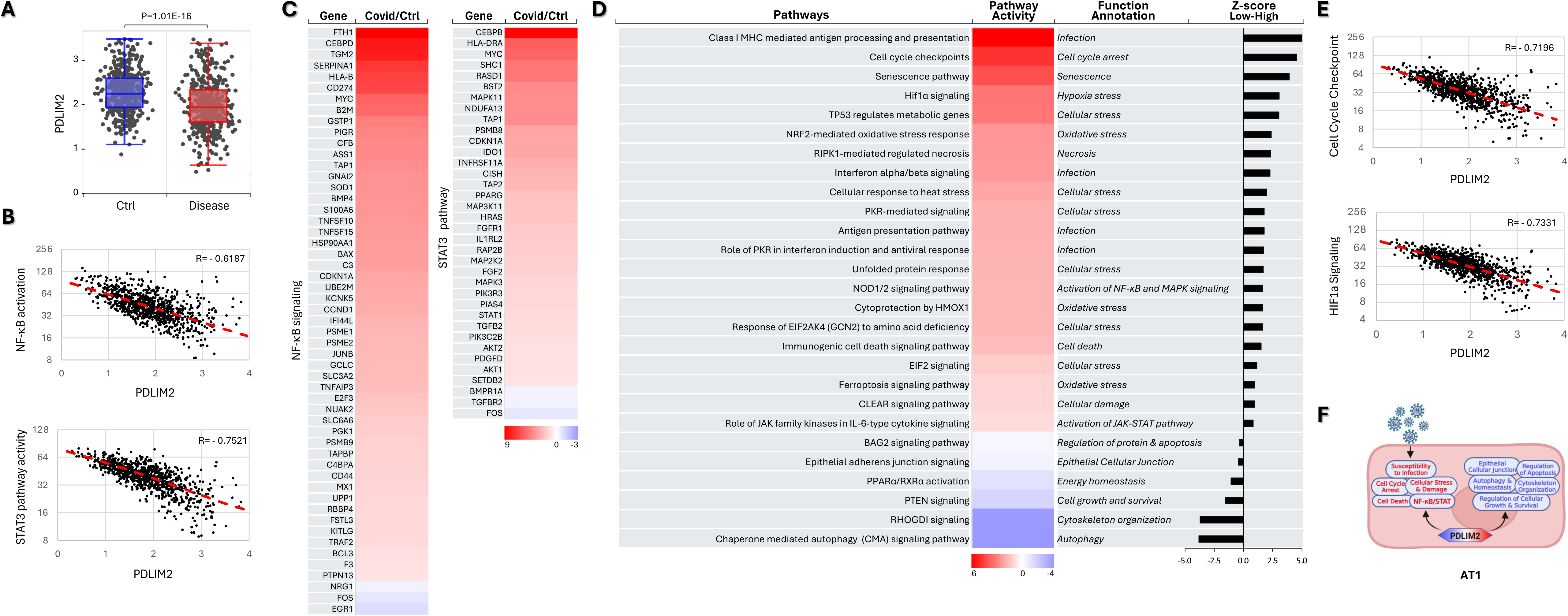
Association of PDLIM2 repression with AT1 cell damage. (A) scRNA-seq data analysis showing PDLIM2 repression in the AT1 cells of COVID-19 patients. P-value was obtained from Student’s *t* test. (B) Gene and pathway correlation showing negative associations of PDLIM2 expression with activations of the NF-κB and STAT3 signaling pathways in human AT1 cells. (C) Heatmap showing differential expression of NF-κB and STAT3 target genes in the AT1 cells of COVID-19 patients in comparison to uninfected ‘healthy’ controls. (D) Heatmap showing differential activation of signaling pathways in PDLIM2-low vs. -high expressing AT1 cells in COVID-19 patients. (E) Gene and pathway correlation showing negative associations of PDLIM2 expression with the activation of the HIF and cell cycle checkpoints signaling pathways in human AT1 cells. (F) Cartoon summary listing the main signaling pathways differentially activated in PDLIM2-low vs. -high expressing AT1 cells in COVID-19 patients.

### Association of PDLIM2 repression with inflammatory activation of airway epithelial cells

Airway epithelial cells (AECs) are the first cells in the lung to encounter inhaled substances like viruses. They form a physical barrier and participate actively in immune responses by releasing chemokines, cytokines, and defensins. However, AECs are also the primary targets for respiratory viruses, and their dysfunction or damage significantly contributes to lung injury and disease progression, including in the context of COVID-19 (Supplementary Table 8 and Supplementary Fig. 7). Mechanistically, we found that PDLIM2 was repressed in the AECs of COVID-19 patients, associating with increased NF-κB, STAT3, cell cycle checkpoints, senescence, clathrin-mediated endocytosis (CME), and pro-inflammatory cytokines but decreased RhoGDI and PPAR signaling pathways (Fig. 8).

**Fig. 8:**
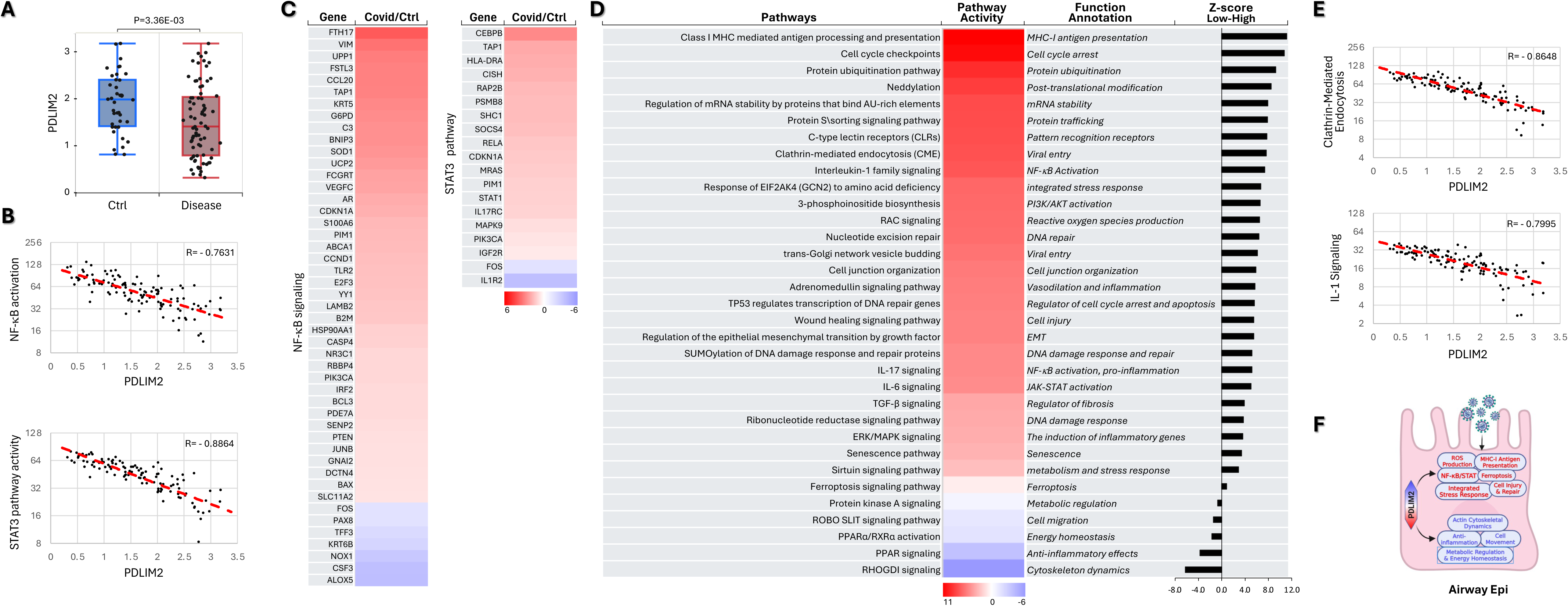
Association of PDLIM2 repression with AEC inflammatory activation. (A) scRNA-seq data analysis showing PDLIM2 repression in the AECs of COVID-19 patients. P-value was obtained from Student’s *t* test. (B) Gene and pathway correlation showing negative associations of PDLIM2 expression with activations of the NF-κB and STAT3 signaling pathways in human ACEs. (C) Heatmap showing differential expression of NF-κB and STAT3 target genes in the AT1 cells of COVID-19 patients in comparison to uninfected controls. (D) Heatmap showing differential activation of signaling pathways in PDLIM2-low vs. -high expressing AECs in COVID-19 patients. (E) Gene and pathway correlation showing negative associations of PDLIM2 expression with the activation of the IL-1 and clathrin-mediated endocytosis signaling pathways in human AECs. (F) Cartoon summary listing the main signaling pathways differentially activated in PDLIM2-low vs. -high expressing AECs in COVID-19 patients.

### Association of PDLIM2 repression with pro-inflammatory and pro-fibrotic activation of lung fibroblasts

Lung fibroblasts, while not being primary targets of lung infections like lung epithelial cells, play a critical role in both lung health and disease. They actively modulate immune responses and can contribute to tissue damage and fibrosis through interactions with other lung cells, particularly epithelial and immune cells. In response to inflammatory signals like those triggered by SARS-CoV-2 infection, they may produce excessive cytokines, chemokines, proteases and extracellular matrices (ECMs) and differentiate into myofibroblasts, potentially leading to pulmonary fibrosis and long-term lung damage (Supplementary Table 9 and Supplementary Fig. 8). The pathological activation of lung fibroblasts in COVID-19 patients involved intrinsic PDLIM2 repression (Fig. 9A), Because the NF-κB, STAT3, cellular stress, pulmonary fibrosis, pathogen-induced cytokine storm. immunogenic cell death and cyclophilin signaling pathways were significantly higher, while the RhoGDI, PPAR and inhibition of matrix metalloproteases (MMPs) signaling pathways were lower in PDLIM2-low lung fibroblasts of COVID-19 patients (Fig. 9B-9F).

**Fig. 9:**
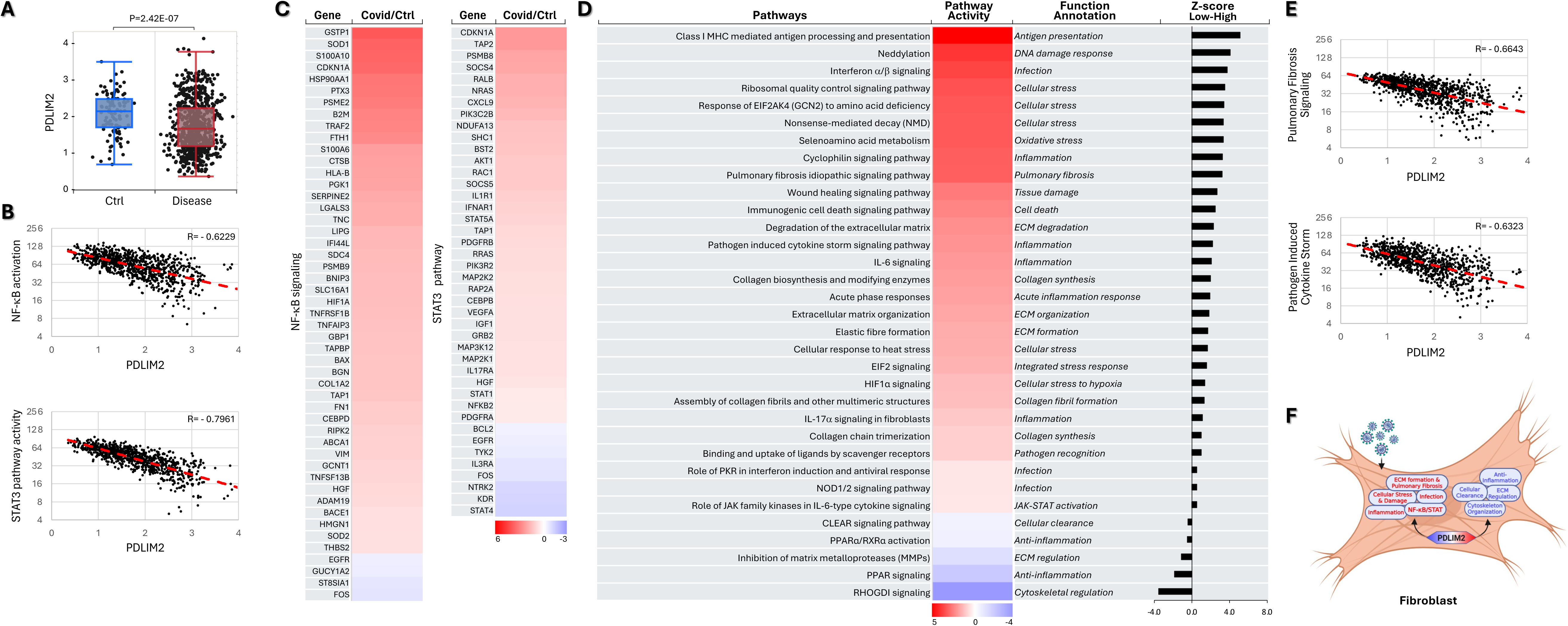
Association of PDLIM2 repression with pro-inflammatory and pro-fibrotic activation of lung fibroblasts. (A) scRNA-seq data analysis showing PDLIM2 repression in the lung fibroblasts of COVID-19 patients. P-value was obtained from Student’s *t* test. (B) Gene and pathway correlation showing negative associations of PDLIM2 expression with activations of the NF-κB and STAT3 signaling pathways in human lung fibroblasts. (C) Heatmap showing differential expression of NF-κB and STAT3 target genes in the lung fibroblasts of COVID-19 patients in comparison to uninfected controls. (D) Heatmap showing differential activation of signaling pathways in PDLIM2-low vs. -high expressing lung fibroblasts in COVID-19 patients. (E) Gene and pathway correlation showing negative associations of PDLIM2 expression with the activation of the pulmonary fibrosis and pathogen-induced cytokine storm signaling pathways in human lung fibroblasts. (F) Cartoon summary listing the main signaling pathways differentially activated in PDLIM2-low vs. -high expressing lung fibroblasts in COVID-19 patients.

## Discussion

Our study highlights both the importance and the clinical applicability of PDLIM2 expression as a biomarker to predict the development and progression of infectious diseases after lung infections by viruses like IV and SARS-CoV-2. The inverse association of PDLIM2 expression in peripheral blood leukocytes with disease severity in respiratory virus-infected humans provides a noninvasive and cost-effective method for diagnosis and prognosis. Furthermore, PDLIM2 repression in BAL cells offers a minimally invasive alternative, while examining PDLIM2 in lung tissues might be a costly and invasive one.

Our study also identifies PDLIM2 repression as a potential new mechanism driving infectious disease onset and progression in virus-infected people, deepening our understanding of viral infection and infectious diseases and revealing new pathophysiological roles of PDLIM2, more importantly, holding significant therapeutic implications. PDLIM2 is repressed in various immune cells and lung structural and functional cells, including lung macrophages, monocytes, neutrophils, T cells, AT1 cells, AT2 cells, AECs, and fibroblasts. Importantly, PDLIM2 repression is associated with aberrant activations of numerous common or cell type-specific signaling pathways, e.g., downregulation of the IL-10, RhoGDI, and PPAR signaling pathways in most cell types, whereas excessive activation of NF-κB, STAT3, cellular stress, senescence, immunogenic cell death, and pathogen-induced cytokine storm signaling pathways in all or most of those cell types, NETosis in neutrophils, and pulmonary fibrosis in fibroblasts. Remarkably, most associates of PDLIM2 are newly identified, with the exceptions of NF-κB and STAT3, which are known downstream targets and mediators of PDLIM2. PDLIM2 likely regulates these new associates directly or indirectly via NF-κB and STAT3, given their frequent crosstalk and shared signaling molecules and target genes. Vice versa, PDLIM2 repression is attributed to oxidative stress-related signaling pathways activated by lung infection, since PDLIM2 expression could be repressed by the reactive oxygen species (ROS)-activated transcription repressor BACH1 [15]. These findings provide mechanistic insights into how PDLIM2 serves as the central nexus of the complex molecular and cellular signaling network crucial for lung health and disease.

Of note, PDLIM2 exhibits multifaceted regulatory effects on signaling pathways, with both positive and negative outcomes depending on cell type and microenvironment. For instance, PDLIM2 repression enhances CXCR4 signaling in monocytes while diminishing it in neutrophils. Similarly, cyclophilin signaling is upregulated in PDLIM2-low neutrophils and fibroblasts but downregulated in PDLIM2-low macrophages. Moreover, while the overall effect of PDLIM2 repression and cell activation in infectious diseases is pathological, they may simultaneously contribute to antiviral activity. Specifically, an increase in antiviral antigen presentation, IFN, and ISGylation signaling pathways is detected across all patient cell types. Their increase in monocytes, AT1 cells, AECs, and fibroblasts involves PDLIM2 repression, whereas PDLIM2 repression attenuates IFN and ISGylation signaling in neutrophils. However, the detected antiviral activity is dominated by the pathological activation of other signaling pathways, especially when the primary target cells of the antiviral activity are defective (e.g., T cells are exhausted, and macrophages are deficient in phagocytosis). In fact, under this situation, excessively high or sustained levels of IFNs mainly contribute to the detrimental inflammatory response, tissue damage, and disease pathogenesis.

Our study has limitations, including its retrospective design, the heterogeneity of the study population, and the lack of detailed information on patients’ demographics and medical history. Additionally, we did not include all cell types, excluding B cells and dendritic cells (DCs) and natural killer (NK) cells, and did not delve into subtypes of cells, particularly for T cells and AECs, which consist of multiple subtypes. Furthermore, the gene expression thresholds were derived internally from this dataset, which may impact generalizability. Despite these limitations, our study presents valuable biomarker discovery, underscoring the need for future prospective studies to validate the prognostic value of PDLIM2 expression in viral infectious diseases, especially influenza and COVID-19.

## Conclusions

In summary, our study demonstrates that low PDLIM2 expression in the context of influenza and COVID-19 correlates with unfavorable molecular and cellular profiles of pro-inflammation and pro-fibrosis and cell injury, and is associated with increased disease severity. These findings establish PDLIM2 as a potential prognostic marker for predicting disease development and progression in individuals infected with viruses like IV and SARS-CoV-2. Furthermore, they highlight the potential of PDLIM2 as a therapeutic target for the prevention and treatment of infectious diseases, especially given the high therapeutic efficacy and safety profile of nanoPDLIM2 in lung cancer models. Considering that PDLIM2 expression is affected in a larger range of cell types in infectious diseases compared to lung cancer, nanoPDLIM2 or other PDLIM2-targeted therapies may offer even better efficacy and safety for patients with infectious diseases.

## Author Contributions

Z.Q. and G.X. conceived and designed the study, led and contributed to all aspects of the analysis, and wrote the manuscript. F.G. searched and analyzed all the data, generated all the figures and tables, and drafted the material and methods section of the manuscript. Y. C assisted data search and analysis. S.D.S provided constructive advice and edited the manuscript. The authors thank other team members for their thoughtful discussion.

## Acknowledgments

This study was financially supported in part by the NIH National Institute of General Medical Sciences (NIGMS) grant R01 GM144890, National Cancer Institute (NCI) grants R01 CA172090 and R01 CA258614, R21 CA259706, National Heart, Lung, and Blood Institute (NHLBI) R01 HL177140, American Cancer Society (ACS) Research Scholar grant RSG-19-166-01-TBG, American Lung Association (ALA) Lung Cancer Discovery Award 821321, and Tobacco Related-Disease Research Program (TRDRP) Research Award T33IR6461.

---

**Fig. S1:**
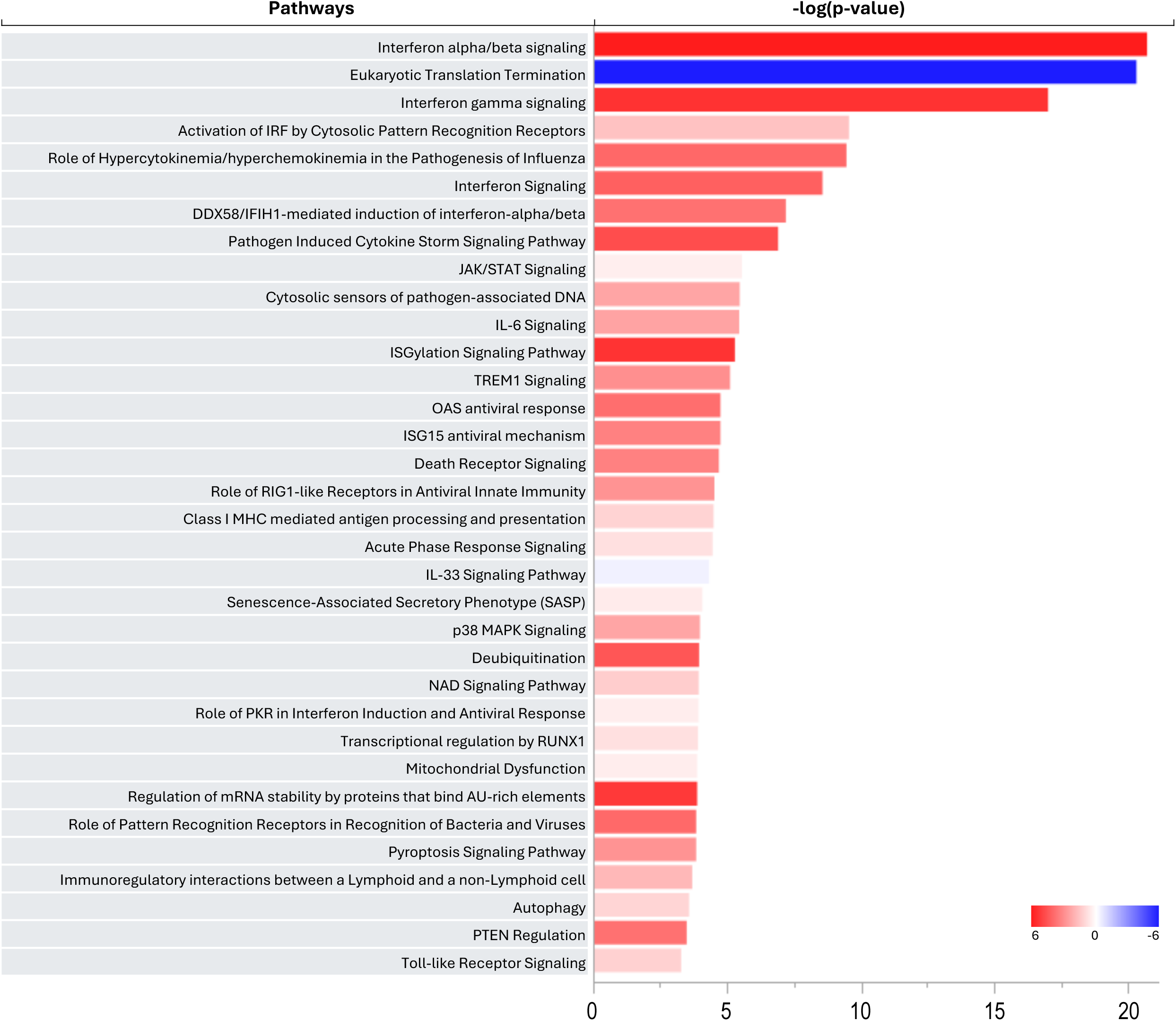
Top pathways altered in patient neutrophils.

**Fig. S2:**
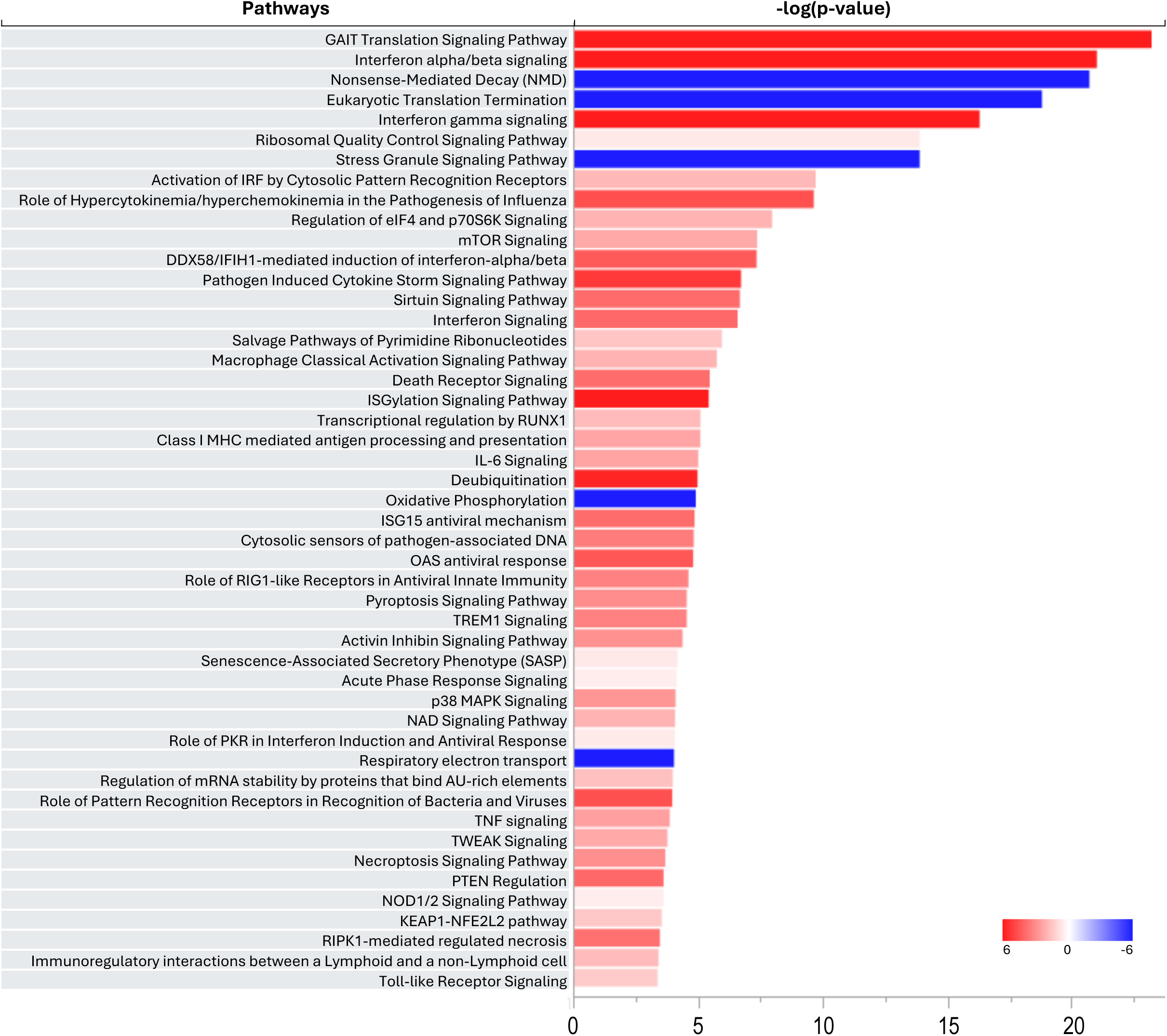
Top pathways altered in patient monocytes.

**Fig. S3:**
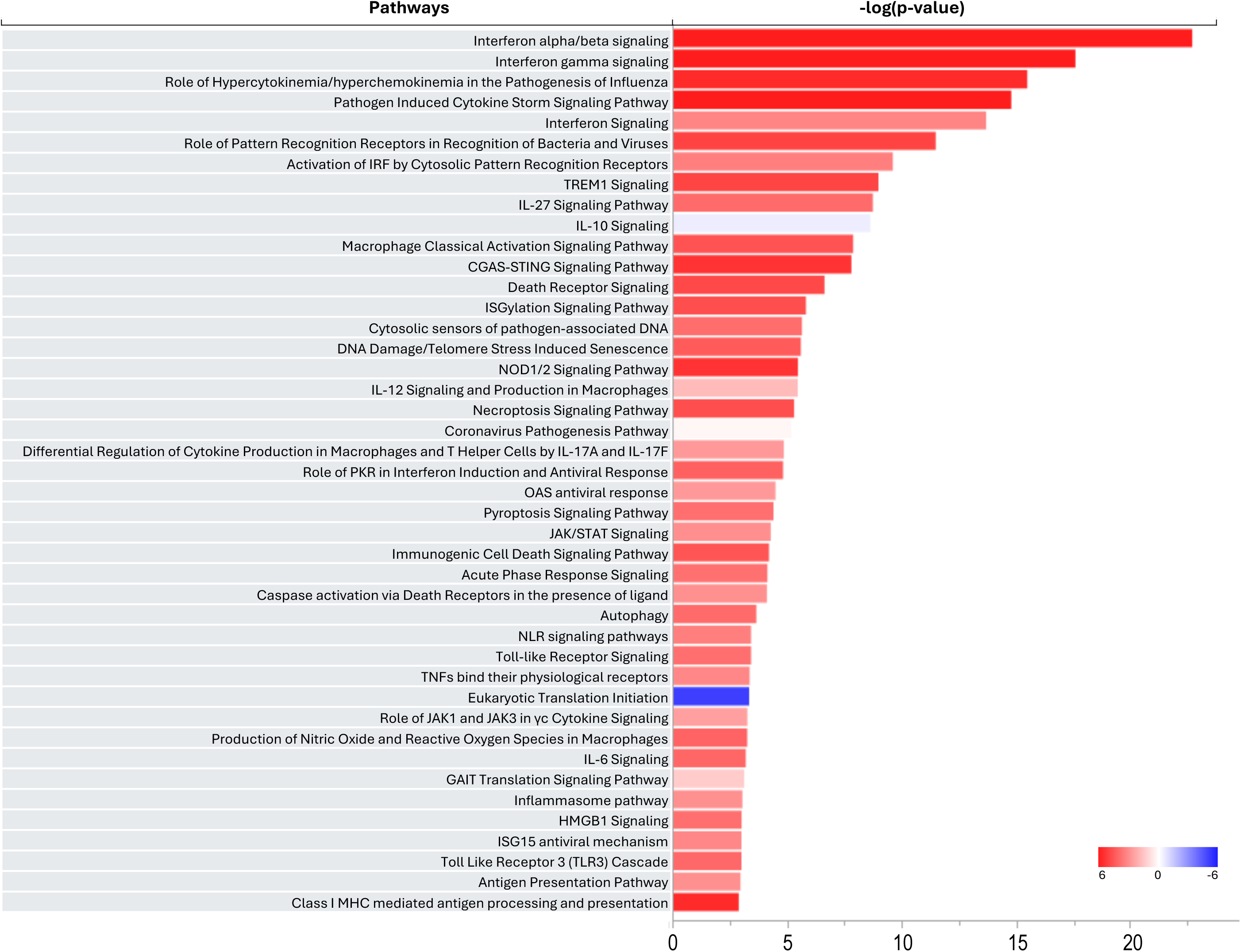
Top pathways altered in patient macrophages.

**Fig. S4:**
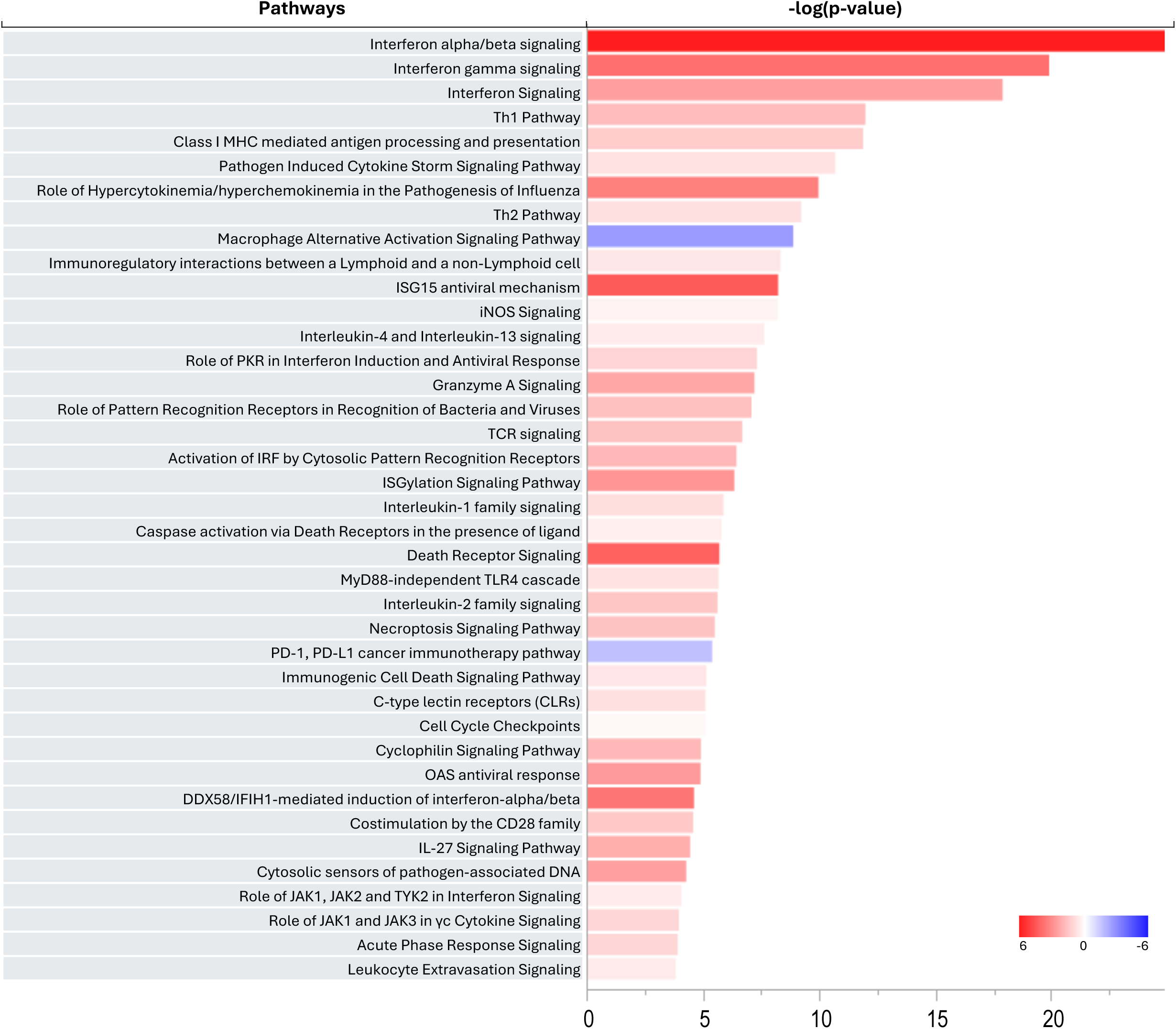
Top pathways altered in patient T cells.

**Fig. S5:**
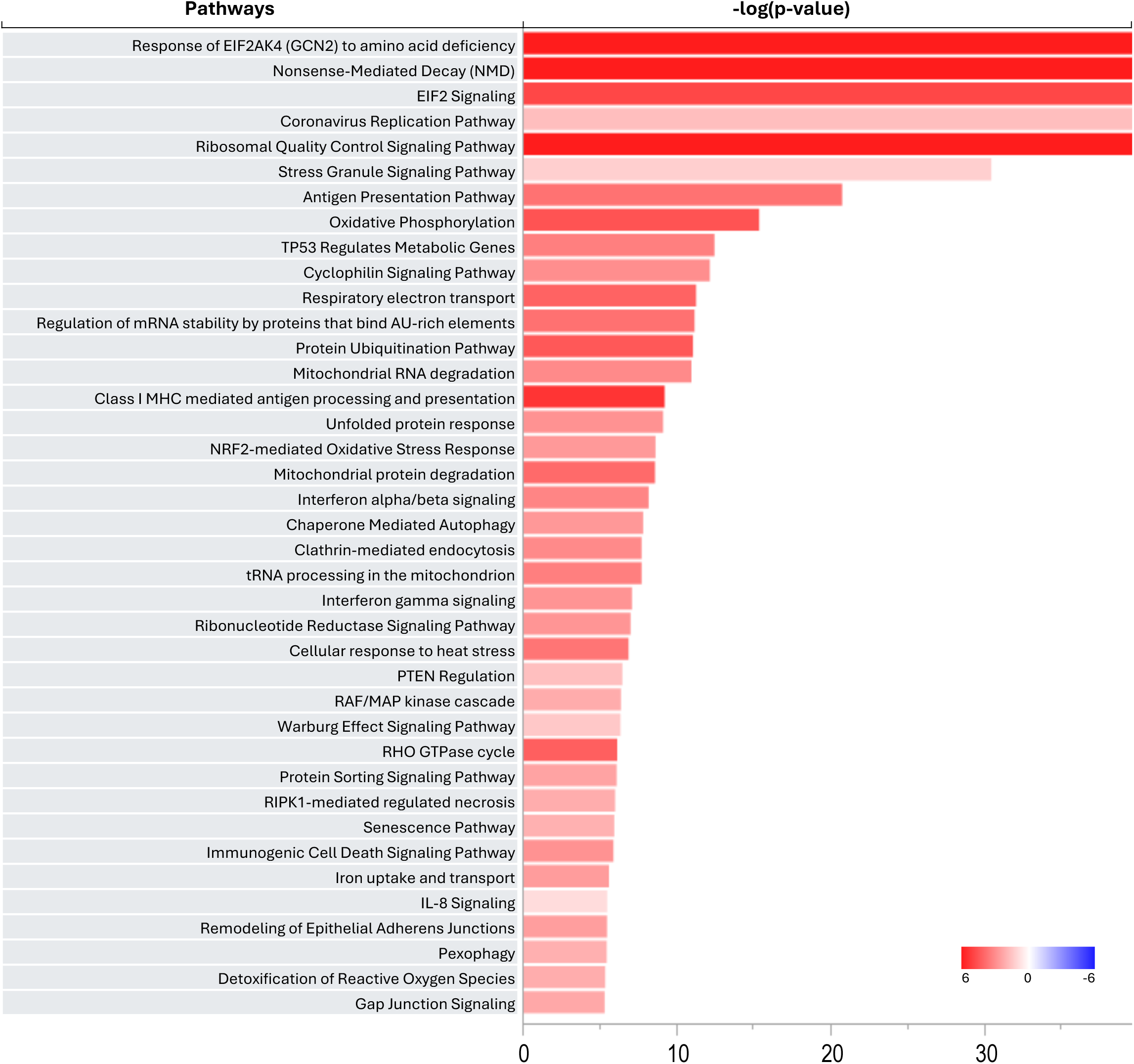
Top pathways altered in patient AT2 cells.

**Fig. S6:**
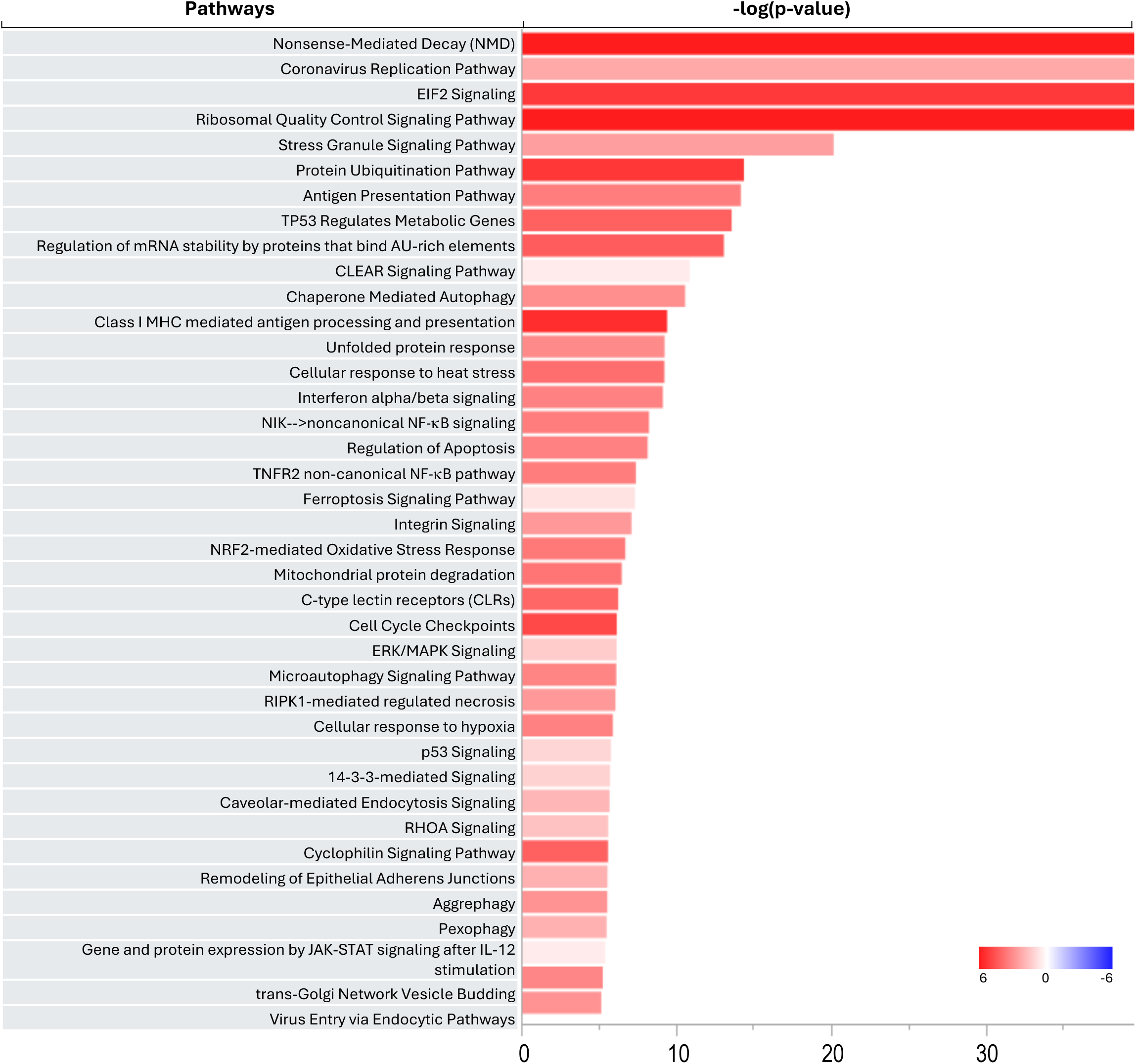
Top pathways altered in patient AT1 cells.

**Fig. S7:**
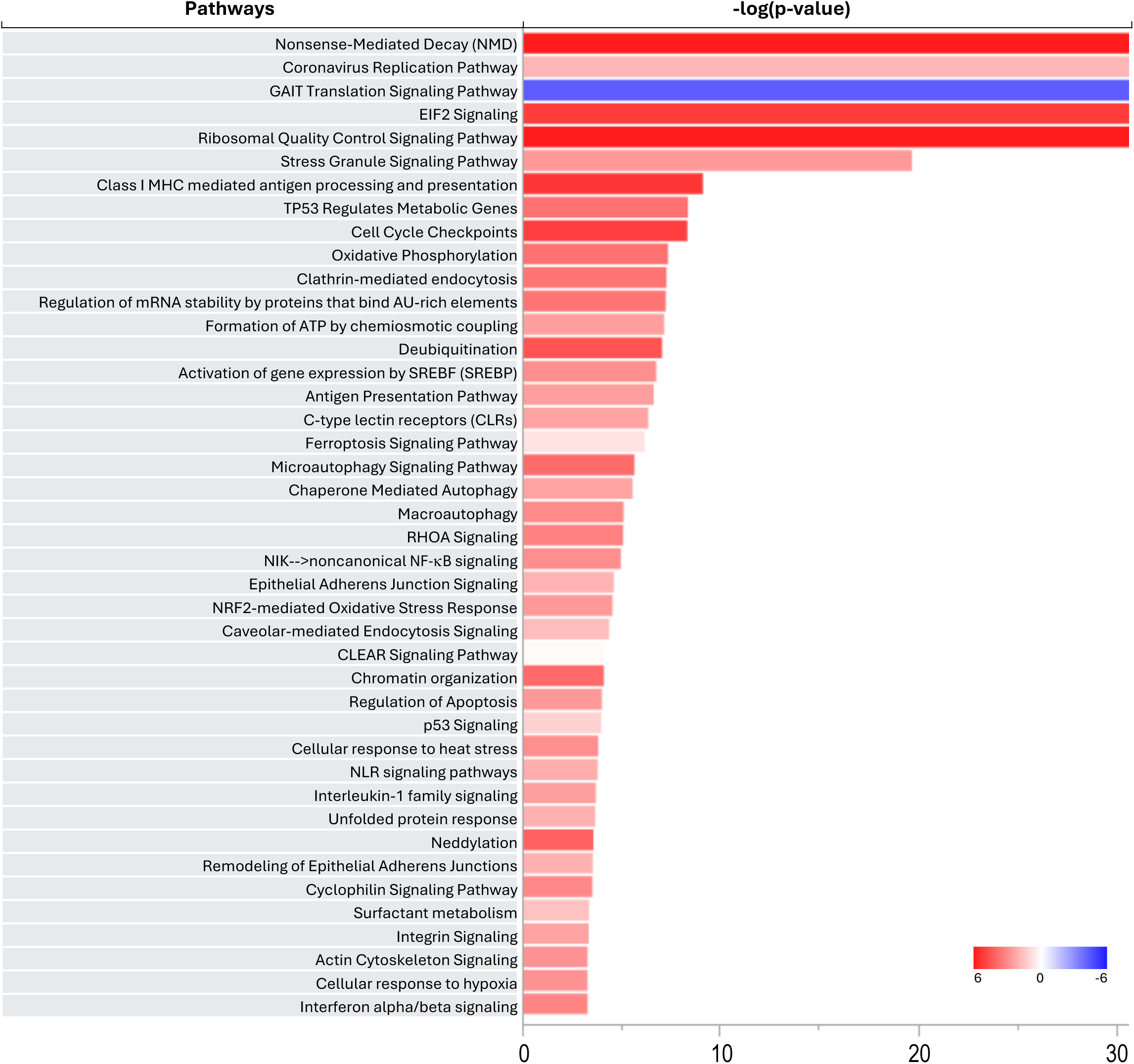
Top pathways altered in patient airway epithelial cells.

**Fig. S8:**
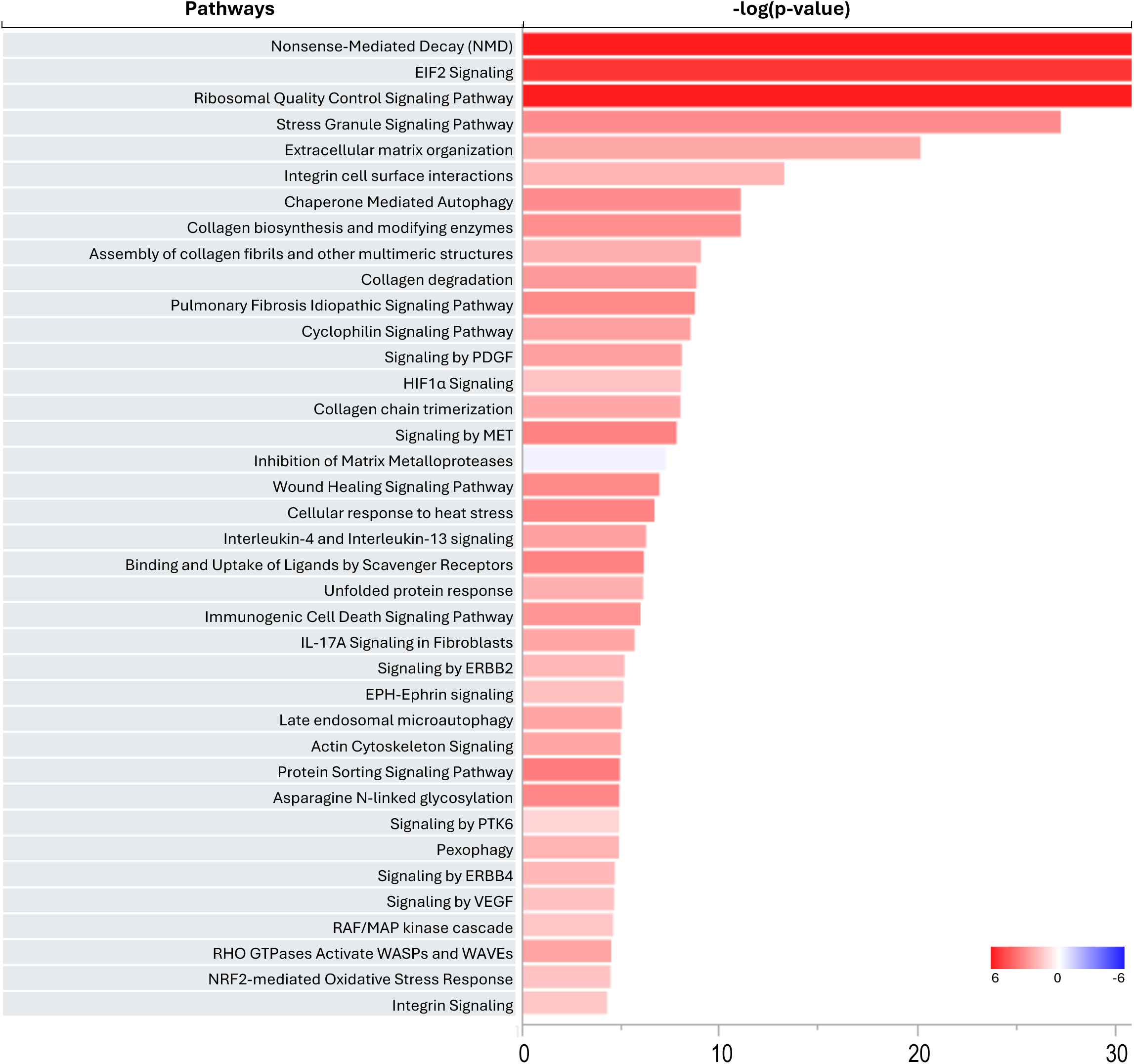
Top pathways altered in patient fibroblasts.

**Table S1:** List of GEO datasets analyzed in the paper.

| GEO | Data | Sample | Virus Type | R scripts on GitHub | Figures in Paper | References |
| --- | --- | --- | --- | --- | --- | --- |
| GSE101702 | Microarray | Blood | Influenza | GSE101702 | Fig.1A | Tang et al., 2019 (PMID:31366921)<br>Zerbib et al., 2020 (PMID: 32066441) |
| GSE157103 | Bulk RNAseq | Blood | SARS-CoV-2 | GSE157103 | Fig.1B | Overmyer et al., 2021 (PMID: 33096026) |
| GSE243629 | scRNAseq | Blood | Influenza | GSE243629 | Fig.2&3 | Zhang et al., 2023 (PMID: 38089584) |
| GSE145926 | scRNAseq | BAL | SARS-CoV-2 | GSE145926 | Fig.4&5 | Liao et al., 2020 (PMID: 32398875)<br>Zhang et al., 2024 (PMID: 39369126) |
| GSE171524 | scRNAseq | Lung Tissue | SARS-CoV-2 | GSE171524 | Fig.6,7,8,9 | Melms et al., 2021 (PMID: 33915568) |

**Table S2:**
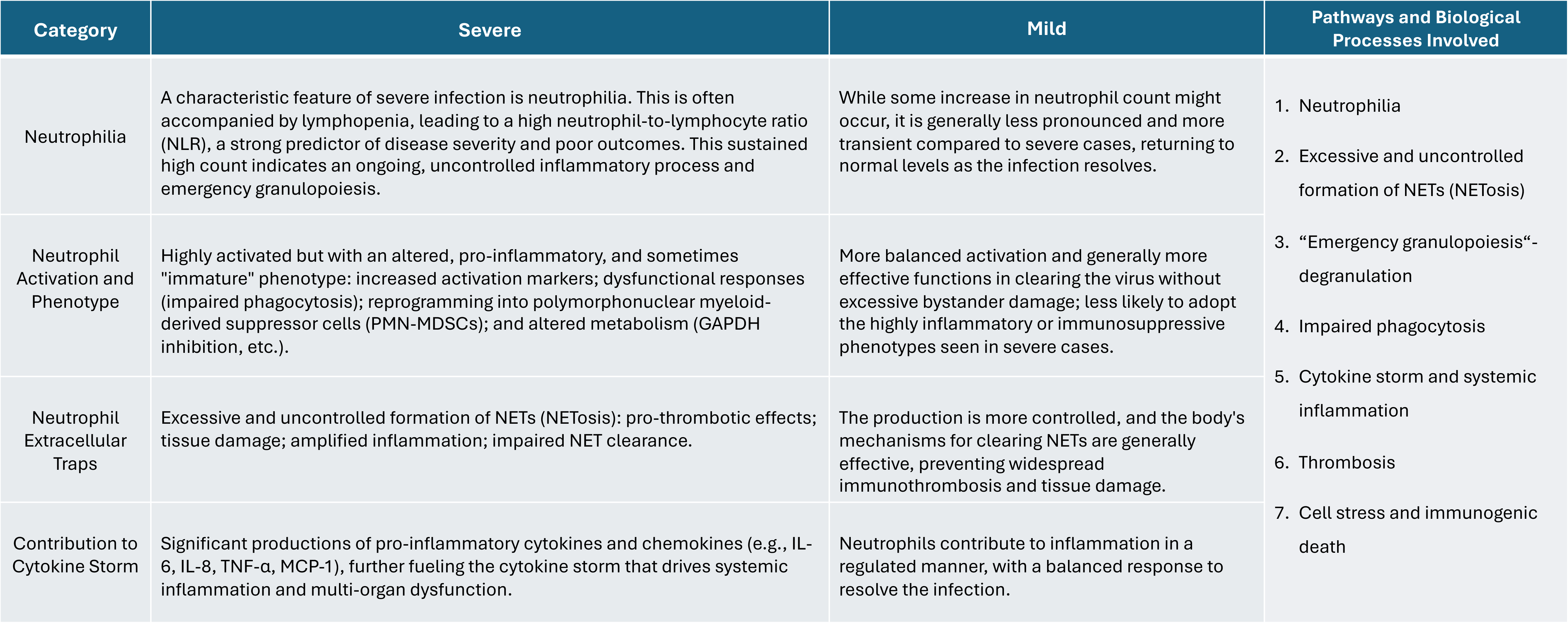
Main differences of neutrophil-related pathways and biological processes in patients with mild and severe conditions by lung viral infection.

**Table S3:** Main differences of monocyte-related pathways and biological processes in patients with mild and severe conditions by lung viral infection.

| Category | Severe | Mild | Pathways and Biological Processes Involved |
| --- | --- | --- | --- |
| Cell numbers and subsets | Often high monocytosis in the peripheral blood, with a particular increase in inflammatory monocytes and lung recruitment. | While monocyte counts may increase during acute infection, the overall increase is typically less pronounced. Monocyte subsets are more balanced, and lung recruitment is more controlled. | 1. Monocytosis<br>2. High monocyte recruitment<br>3. Inflammasome activation<br>4. Hyper-inflammation<br>5. Cell stress and immunogenic cell death<br>6. Increased antigen presentation |
| Activation and pro-inflammatory cytokine production | High production of pro-inflammatory cytokines and chemokines: $\text{TNF}\alpha$, IL-6, type I IFNs, MCP-1 (CCL2), MIP-1α (CCL3), and IP-10 (CXCL10); and hyper inflammasome activation (e.g., NLRP3 activation for IL-1 and IL-18 release) and pyroptosis. | More regulated and self-limiting. The cytokine production is proportionate to the viral load and contributes to effective viral clearance without causing excessive systemic inflammation. | |
| Antigen presentation | Downregulation of HLA-DR leading to impaired antigen presentation and T cell activation (immunoparalysis) | Normal or even upregulated HLA-DR expression |  |
| Differentiation and role in lung pathology | The inflammatory monocytes recruited to the lungs differentiate into highly pro-inflammatory macrophages and dendritic cells. These cells contribute significantly to lung injury and ARDS by: direct tissue damage; exacerbating inflammation; promoting fibrosis; secondary infections. | Monocytes effectively differentiate into macrophages and dendritic cells that contribute to viral clearance and then change to a reparative phenotype, facilitating tissue healing and inflammation resolution. |  |

**Table S4:** Main differences of macrophage-related pathways and biological processes in patients with mild and severe conditions by lung viral infection.

| Category | Severe | Mild | Pathways and Biological Processes Involved |
| --- | --- | --- | --- |
| Macrophage activation and phenotype (polarization) | Hyper pro-inflammatory M1 polarization and state; increased inflammasome activation (e.g., NLRP3 inflammasome); impaired inflammation resolution activity; dysfunctional efferocytosis; and altered surface markers (increased CD163). | Controlled inflammation; effective viral clearance; and balanced polarization. | 1. M1/M2 macrophage<br>2. Phagocytosis and efferocytosis<br>3. Cytokine storm<br>4. Production of nitric oxide and reactive oxygen species in macrophages<br>5. Inflammasome activation<br>6. The systemic inflammatory response syndrome (SIRS)<br>7. The increased inflammatory monocyte-derived macrophages particularly in the lung<br>8. Cell death |
| Cytokine production | Profound and sustained production of pro-inflammatory cytokines and chemokines (e.g., IL-1β, IL-6, TNF-α, MCP-1, MIP-1α, IP-10, etc.), leading to the systemic inflammatory response syndrome (SIRS) and potential multi-organ dysfunction. | More transient and localized production of cytokines, sufficient to control the infection without causing widespread hyper-inflammation. | |
| Recruitment and accumulation | Increased recruitment and accumulation of inflammatory monocytes and differentiation into macrophages, particularly in the lung. | While macrophages are recruited to the infection sites, their numbers and activity are more balanced. |  |
| Summary | The dysregulation and sustained pro-inflammatory activation of macrophages lead to a runaway inflammatory process that damages host tissues and is a major driver of pathology | A controlled, immune response that resolves inflammation after viral clearance |  |

**Table S5:** Main differences of T-cell-related pathways and biological processes in patients with mild and severe conditions by lung viral infection.

| Category | Severe | Mild | Pathways and Biological Processes Involved |
| --- | --- | --- | --- |
| Lymphopenia | A hallmark of severe infectious diseases like COVID-19 is pronounced lymphopenia, particularly CD4+ helper and CD8+ cytotoxic T cells: redistribution/sequestration; apoptosis/pyroptosis; bone marrow suppression; and T cell exhaustion. | While some degree of transient lymphopenia may occur, it is generally less severe, and T-cell counts tend to recover more quickly compared to severe cases. | 1. Lymphopenia<br>2. T cell exhaustion<br>3. Hyperinflammation<br>4. Cytokine storm<br>5. Cell stress<br>6. Senescence<br>7. Apoptosis/pyroptosis |
| T cell exhaustion | Functionally impaired due to prolonged and intense antigen stimulation and exposure to high levels of inflammatory cytokines | T cells generally maintain better functionality and are less prone to severe exhaustion. |  |
| Virus-specific T cell response | Patients with severe COVID-19 may show a robust and sometimes even stronger sars-cov-2-specific T cell response in terms of magnitude. However, the response is ineffective due to lymphopenia and T-cell exhaustion, leading to hyperinflammation rather than effective viral clearance. | Patients typically develop a more effective and well-coordinated sars-cov-2-specific T cell response. They often have better memory T-cell responses that contribute to long-term immunity. |  |
| Cytokine production | The overall production of key antiviral cytokines like IFN-γ by T cells might be lower, however, many other inflammatory cytokines are upregulated. | T cells produce appropriate levels of antiviral cytokines, particularly IFN-γ | |
| T cell subsets | Expansion of highly differentiated T cells (e.g., terminal effector memory T cells re-expressing CD45RA), a sign of chronic immune activation and exhaustion. | A more balanced T cell subset distribution, with effective memory T cell populations that contribute to rapid responses upon re-exposure. |  |

**Table S6:** Main differences of AT2 cell-related pathways and biological processes in patients with mild and severe conditions by lung viral infection.

| Category | Severe | Mild | Pathways and biological processes involved |
| --- | --- | --- | --- |
| Viral infection and replication | Extensive and sustained virus infection and replication within AT2 cells, leading to direct viral cytotoxicity, high viral load (persistent inflammation), senescence-associated secretory phenotype (SASP), and pro-inflammatory phenotype (cytokine storm and chronic inflammation). | While AT2 cells are infected, the viral load is generally lower, and the infection is more effectively controlled by the early immune response. The extent of direct viral damage and cell death is less widespread, allowing for more effective recovery. | 1. Higher infection<br>2. Hyperinflammation, cytokine storm<br>3. Cell death and senescence<br>4. Surfactant dysfunction<br>5. Compromised antiviral interferon response<br>6. Impaired regeneration (proliferation and differentiation)<br>7. Fibrosis promotion |
| Surfactant production | Reduced surfactant production, surfactant dysfunction, and alveolar collapse (atelectasis). | Surfactant production may be transiently affected, no widespread surfactant deficiency and subsequent alveolar collapse. |  |
| Role in lung repair and regeneration | Impaired proliferation and differentiation, aberrant differentiation/maladaptive repair, and fibrosis promotion. | Effective proliferation and differentiation, resolution of inflammation |  |
| Immunomodulatory role | Dysfunctional AT2 cells contribute to the cytokine storm by releasing pro-inflammatory mediators. Their impaired function can also lead to a less effective early antiviral interferon response, potentially allowing for more viral replication and exacerbating the immune-mediated damage | AT2 cells contribute to a more balanced innate immune response, producing interferons and other mediators that help control viral replication without triggering overwhelming inflammation. |  |

**Table S7:** Main differences of AT1 cell-related pathways and biological processes in patients with mild and severe conditions by lung viral infection.

| Category | Severe | Mild | Pathways and biological processes involved |
| --- | --- | --- | --- |
| Direct damage and cell death | High damage and death (pyroptosis and apoptosis) by the overwhelming infection and inflammatory response, including cytokine storm, neutrophil extracellular traps (nets), oxidative stress; and hyaline membrane formation. | The damage is generally localized and less widespread. The inflammatory response is more controlled, limiting bystander injury to AT1 cells. | 1. Viral infection<br>2. High production of inflammation cytokines and chemokines/Cytokine storm<br>3. Cellular stress: oxidative, inflammation, hypoxemia, etc.<br>4. Cell damage and death: apoptosis, pyroptosis, etc.<br>5. Pro-fibrotic environment<br>6. ARDS<br>7. Excessive and uncontrolled recruitment of inflammatory cells |
| Impaired repair and regeneration | Defects in AT1 cell differentiation from infected or highly stressed AT2 cells or pause at the “transitional state” of AT2 to AT1 cells. | AT2 cells effectively proliferate and differentiate into functional AT1 cells, ensuring efficient repair of the alveolar epithelium. |  |
| Functional consequences | Severe impairment of gas exchange; acute respiratory distress syndrome (ARDS); persistent hypoxemia; and long-term lung damage. | Minimal impact on gas exchange due to limited and transient AT1 cell damage and effective repair. |  |
| Immunomodulatory role | Sustained release of cytokines and chemokines; excessive and uncontrolled recruitment of inflammatory cells; and widespread death and the loss of the epithelial barrier. | Controlled and localized immune response from their cytokine and chemokine production; and the integrity of AT1 layer acting as a barrier for pathogens and immune cells. |  |

**Table S8:** Main differences of AEC-related pathways and biological processes in patients with mild and severe conditions by lung viral infection.

| Category | Severe | Mild | Pathways and Biological Processes Involved |
| --- | --- | --- | --- |
| Viral infection and spread | Extensive viral infection and deep spread; and high viral loads | The initial infection might be more localized, often starting in the upper respiratory tract (nasal mucosa). | 1. Viral infection and spread<br>2. Hyper inflammatory, exacerbated/systemic cytokine/chemokine storm<br>3. Class I MHC mediated antigen processing and presentation<br>4. IFN response, pattern recognition<br>5. Immune cell infiltration<br>6. Severe cell stress, damage, injury, death<br>7. Impaired regeneration/repair |
| Antiviral response | Dysregulated antiviral response; delayed/suppressed IFN response; hyperinflammation; immune cell infiltration; and bystander damage. | More effective and controlled early antiviral IFN response with timely IFN production; controlled pro-inflammatory cytokines and chemokines release |  |
| Cellular damage and repair | Severe cellular damage and impaired repair; widespread cell injury and death; hyaline membrane formation; loss of barrier integrity; disrupted mucociliary clearance; and impaired regeneration and fibrosis | Transient cilia shedding and mucociliary dysfunction; limited programmed cell death (necroptosis); preserved epithelial repair and regeneration |  |
| Immunomodulatory roles | Impaired pattern recognition and early warning; immunomodulatory role becomes maladaptive and contributes to pathogenesis: delayed or dysregulated immune response, exacerbated cytokine/chemokine storm. | Effective pattern recognition and early warning; and controlled and proportionate production of IFNs and pro-inflammatory cytokines/chemokines, restricting viral replication and shaping immunity . |  |

**Table S9:**
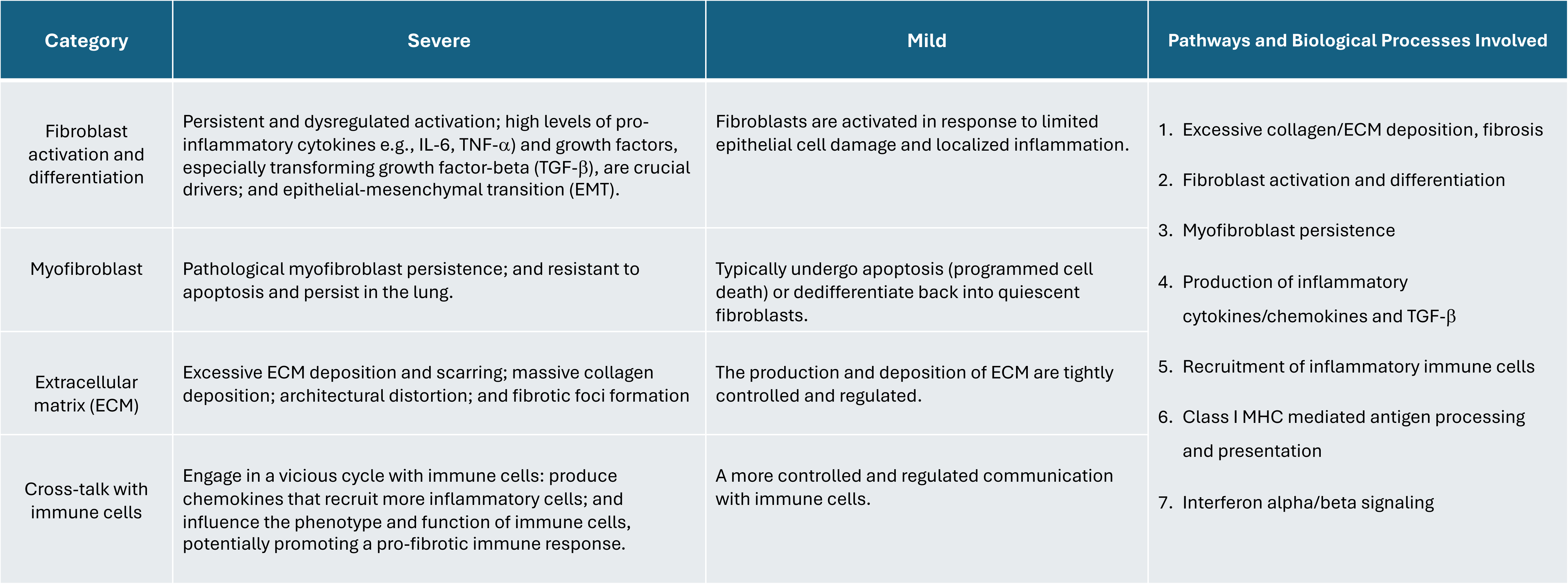
Main differences of lung fibroblast-related pathways and biological processes in patients with mild and severe conditions by lung viral infection.

## References

1. Prabhu FR, Sikes AR, Sulapas I. Pulmonary infections. Family Medicine. 2016 Feb 17:1083–101. doi: 10.1007/978-3-319-04414-9_91. PMCID: PMC7120011.

2. Niederman MS, Torres A. Respiratory infections. Eur Respir Rev 2022; 31: 220150 [DOI: 10.1183/16000617.0150-2022].

3. Weissferdt A. Infectious lung disease. Diagnostic Thoracic Pathology. 2019 Dec 3:3–71. doi: 10.1007/978-3-030-36438-0_1. PMCID: PMC7335808.

4. Rolfo C, Meshulami N, Russo A, Krammer F, García-Sastre A, Mack PC, Gomez JE, Bhardwaj N, Benyounes A, Sirera R, Moore A, Rohs N, Henschke CI, Yankelevitz D, King J, Shyr Y, Bunn PA Jr, Minna JD, Hirsch FR. Lung cancer and severe acute respiratory syndrome coronavirus 2 infection: identifying important knowledge gaps for Investigation. J Thorac Oncol. 2022 Feb;17(2):214–227. doi: 10.1016/j.jtho.2021.11.001. Epub 2021 Nov 10. PMID: 34774792; PMCID: PMC8579698.

5. Kinslow CJ, Wang Y, Liu Y, Zuev KM, Chaudhary KR, Wang TJC, Donalek C, Amori M, Cheng SK. Influenza activity and regional mortality for non-small cell lung cancer. Sci Rep. 2023 Dec 7;13(1):21674. doi: 10.1038/s41598-023-47173-x. PMID: 38065996; PMCID: PMC10709588.

6. Suri C, Pande B, Sahithi LS, Sahu T, Verma HK. Interplay between lung diseases and viral infections: a comprehensive review. Microorganisms. 2024 Oct 8;12(10):2030. doi: 10.3390/microorganisms12102030. PMID: 39458339; PMCID: PMC11510474.

7. Sun, F., Li, L., Yan, P., Zhou, J., Shapiro, S. D., Xiao, G. and Qu, Z. (2019). Causative role of PDLIM2 epigenetic repression in lung cancer and therapeutic resistance. Nat Commun. 10(1): 5324. 10.1038/s41467-019-13331-x

8. Sun, F., Yan, P., Xiao, Y., Zhang, H., Shapiro, S. D., Xiao, G. and Qu, Z. (2024). Improving PD-1 blockade plus chemotherapy for complete remission of lung cancer by nanoPDLIM2. eLife. 12: RP89638. 10.7554/eLife.89638.3

9. Ashouri K, Krause H, Eilliott A, Liu SV, Ma PC, Halmos B, Qu Z, Xiao G, Vanderwalde A, Nieva JJ. Characterization of PDLIM2 in non-small cell lung cancer. Cancer Res. 2024; 84(S6): 5201. 10.1158/1538-7445.AM2024-5201

10. Le TH, Sun F, Xiao G, Qu Z. NanoPDLIM2-based combination therapy for lung cancer treatment in mouse preclinical studies. Bio-protocol. 2025;in press. DOI: 10.21769/BioProtoc.5437

11. Gao F, Liu X, Sun F, Xiao Y, Xiao G, Qu Z. Myeloid PDLIM2 repression as a common mechanism of infection susceptibility in lung diseases. 2025; in review.

12. Torrado M, Senatorov VV, Trivedi R, Fariss RN, Tomarev SI. Pdlim2, a novel PDZ-LIM domain protein, interacts with a-actinins and filamin A. Invest Ophthalmol Vis Sci. 2004 Nov;45(11):3955–63. doi: 10.1167/iovs.04-0721.

13. Loughran, G., Healy, N. C., Kiely, P. A., Huigsloot, M., Kedersha, N. L. and O’Connor, R. (2005). Mystique is a new insulin-like growth factor-I-regulated PDZ-LIM domain protein that promotes cell attachment and migration and suppresses Anchorage-independent growth. Mol Biol Cell. 16(4): 1811–1822. 10.1091/mbc.e04-12-1052

14. Tanaka T, Soriano MA, Grusby MJ. SLIM is a nuclear ubiquitin E3 ligase that negatively regulates STAT signaling. Immunity. 2005 Jun;22(6):729–36. doi: 10.1016/j.immuni.2005.04.008.

15. Li, L., Sun, F., Han, L., Liu, X., Xiao, Y., Gregory, A. D., Shapiro, S. D., Xiao, G. and Qu, Z. (2021). PDLIM2 repression by ROS in alveolar macrophages promotes lung tumorigenesis. JCI Insight. 6(5): e144394. 10.1172/jci.insight.144394

16. Guo, Z. S. and Qu, Z. (2021). PDLIM2: Signaling pathways and functions in cancer suppression and host immunity. Biochim Biophys Acta Rev Cancer. 1876(2): 188630. 10.1016/j.bbcan.2021.188630

17. Yan P, Fu J, Qu Z, Li S, Tanaka T, Grusby MJ, Xiao G. PDLIM2 suppresses human T-cell leukemia virus type I Tax-mediated tumorigenesis by targeting Tax into the nuclear matrix for proteasomal degradation. Blood. 2009;113:4370–4380.

18. Yan P, Qu Z, Ishikawa C, Mori N, Xiao G. Human T-cell leukemia virus type I-mediated repression of PDZ-LIM domain-containing protein 2 involves DNA methylation but independent of the viral oncoprotein Tax. Neoplasia. 2009;11:1036–1041.

19. Fu J, Yan P, Li S, Qu Z, Xiao G. Molecular determinants of PDLIM2 in suppressing HTLV-I Tax-mediated tumorigenesis. Oncogene. 2010;29:6499–6507.

20. Qu Z, Xiao G. Human T-cell lymphotropic virus: a model of NF-κB associated tumorigenesis. Viruses. 2011; 3(6):714–749.

21. Qu, Z., Fu, J., Yan, P., Hu, J., Cheng, S. Y. and Xiao, G. (2010). Epigenetic repression of PDZ-LIM domain-containing protein 2: implications for the biology and treatment of breast cancer. J Biol Chem. 285(16): 11786–11792. 10.1074/jbc.M109.086561

22. Qu, Z., Yan, P., Fu, J., Jiang, J., Grusby, M. J., Smithgall, T. E. and Xiao, G. (2010). DNA methylation-dependent repression of PDZ-LIM domain-containing protein 2 in colon cancer and its role as a potential therapeutic target. Cancer Res. (70 (5): 1766–1772. *Cancer Res.* 70: 1766–1772. 10.1158/0008-5472.CAN-09-3263

23. Sun, F., Xiao, Y. and Qu, Z. (2015). Oncovirus Kaposi sarcoma herpesvirus (KSHV) represses tumor suppressor PDLIM2 to persistently activate nuclear factor κB (NF-κB) and STAT3 transcription factors for tumorigenesis and tumor maintenance. J Biol Chem. 290(12): 7362–7368. 10.1074/jbc.C115.637918.

24. Vanoirbeek E, Eelen G, Verlinden L, Carmeliet G, Mathieu C, Bouillon R, O’Connor R, Xiao G, Verstuyf A. PDLIM2 expression is driven by vitamin D and is involved in the pro-adhesion, and anti-migration and -invasion activity of vitamin D. Oncogene. 2014;33(15):1904–1911.

25. Qu, Z., Fu, J., Ma, H., Zhou, J., Jin, M., Mapara, M. Y., Grusby, M. J. and Xiao, G. (2012). PDLIM2 restricts Th1 and Th17 differentiation and prevents autoimmune disease. Cell Biosci. 2(1): 23. 10.1186/2045-3701-2-23

26. Tanaka, T., Grusby, M. J. and Kaisho, T. (2007). PDLIM2-mediated termination of transcription factor NF-κB activation by intranuclear sequestration and degradation of the p65 subunit. Nat Immunol. 8(6): 584–591. 10.1038/ni1464

27. Xiao G, Rabson AB, Young W, Qing G, Qu Z. Alternative pathways of NF-κB activation: a double-edged sword in health and disease. Cytokine Growth Factor Rev. 2006;17:281–293.

28. Xiao, G. and Fu, J. (2011). NF-κB and cancer: a paradigm of Yin-Yang. Am J Cancer Res. 1(2): 192–221. https://pmc.ncbi.nlm.nih.gov/articles/PMC3180046

29. Li, L., Han, L., Sun, F., Zhou, J., Ohaegbulam, K. C., Tang, X., Zang, X., Steinbrecher, K. A., Qu, Z. and Xiao, G. (2018). NF-κB RelA renders tumor-associated macrophages resistant to and capable of directly suppressing CD8(+) T cells for tumor promotion. Oncoimmunology. 7(6): e1435250. 10.1080/2162402X.2018.1435250

30. Li L, Han L, Qu Z. NF-κB RelA is a cell-intrinsic metabolic checkpoint restricting glycolysis. Cell Biosci. 2024 Jan 20;14(1):11. doi: 10.1186/s13578-024-01196-7.

31. Qing G, Qu Z, Xiao G. Endoproteolytic processing of C-terminally truncated NF-κB2 precursors at κB-containing promoters. Proc Natl Acad Sci U S A. 2007 Mar 27;104(13):5324–9. doi: 10.1073/pnas.0609914104.

32. Sun, F., Qu, Z., Xiao, Y., Zhou, J., Burns, T. F., Stabile, L. P., Siegfried, J. M. and Xiao, G. (2016). NF-κB1 p105 suppresses lung tumorigenesis through the Tpl2 kinase but independently of its NF-κB function. Oncogene. 35(18): 2299–2310. 10.1038/onc.2015.299

33. Sun F, Xiao Y, Shapiro SD, Qu Z, Xiao G. Critical and distinct roles of cell type-specific NF-κB2 in lung cancer. JCI Insight. 2024 Feb 22;9(4):e164188. doi: 10.1172/jci.insight.164188.

34. Fu J, Qu Z, Yan P, Ishikawa C, Aqeilan RI, Rabson AB, Xiao G. The tumor suppressor gene WWOX links the canonical and noncanonical NF-κB pathways in HTLV-I Tax-mediated tumorigenesis. Blood. 2011 Feb 3;117(5):1652–61. doi: 10.1182/blood-2010-08-303073.

35. Yan P, Qing G, Qu Z, Wu CC, Rabson A, Xiao G. Targeting autophagic regulation of NF-κB activation in HTLV-I transformed cells by geldanamycin: Implications for therapeutic interventions. Autophagy. 2007;3(6):600–603.

36. Zhou, J., Qu, Z., Yan, S., Sun, F., Whitsett, J. A., Shapiro, S. D. and Xiao, G. (2015). Differential roles of STAT3 in the initiation and growth of lung cancer. Oncogene. 34 (29): 3804–3814. 10.1038/onc.2014.318

37. Qu, Z., Sun, F., Zhou, J., Li, L., Shapiro, S. D. and Xiao, G. (2015). Interleukin-6 prevents the initiation but enhances the progression of lung cancer. Cancer Res. 75(16): 3209–3215. 10.1158/0008-5472.CAN-14-3042

38. Zhou, J., Qu, Z., Sun, F., Han, L., Li, L., Yan, S., Stabile, L. P., Chen, L. F., Siegfried, J. M. and Xiao, G. (2017) Myeloid STAT3 promotes lung tumorigenesis by transforming tumor immunosurveillance into tumor-promoting inflammation. Cancer Immunol Res. 5(3): 257–268. 10.1158/2326-6066.CIR-16-0073

39. Zhang H, Xiao Y, Gao F, Le TH, Shapiro SD, Xiao G, Qu Z. Alveolar epithelial NF-κB/RelA guards the lung against bacterial infection. 2025;in review.

40. Sun F, Xiao G, Qu Z. PDLIM2 is a novel E5 ubiquitin ligase enhancer that stabilizes ROC1 and recruits the ROC1-SCF ubiquitin ligase to ubiquitinate and degrade NF-κB RelA. Cell Biosci. 2024;14(1):99.

41. Qu Z, Qing G, Rabson A, Xiao G. Tax deregulation of NF-κB2 p100 processing involves both β-TrCP-dependent and -independent mechanisms. J Biol Chem. 2004 Oct 22;279(43):44563–72. doi: 10.1074/jbc.M403689200.

42. Tang BM, Shojaei M, Teoh S, Meyers A, Ho J, Ball TB, Keynan Y, Pisipati A, Kumar A, Eisen DP, Lai K, Gillett M, Santram R, Geffers R, Schreiber J, Mozhui K, Huang S, Parnell GP, Nalos M, Holubova M, Chew T, Booth D, Kumar A, McLean A, Schughart K. Neutrophils-related host factors associated with severe disease and fatality in patients with influenza infection. Nat Commun. 2019 Jul 31;10(1):3422. doi: 10.1038/s41467-019-11249-y. PMID: 31366921; PMCID: PMC6668409.

43. Zerbib Y, Jenkins EK, Shojaei M, Meyers AFA, Ho J, Ball TB, Keynan Y, Pisipati A, Kumar A, Kumar A, Nalos M, Tang BM, Schughart K, McLean A; Nepean Genomic Research Group. Pathway mapping of leukocyte transcriptome in influenza patients reveals distinct pathogenic mechanisms associated with progression to severe infection. BMC Med Genomics. 2020 Feb 17;13(1):28. doi: 10.1186/s12920-020-0672-7. PMID: 32066441; PMCID: PMC7027223.

44. Overmyer KA, Shishkova E, Miller IJ, Balnis J, Bernstein MN, Peters-Clarke TM, Meyer JG, Quan Q, Muehlbauer LK, Trujillo EA, He Y, Chopra A, Chieng HC, Tiwari A, Judson MA, Paulson B, Brademan DR, Zhu Y, Serrano LR, Linke V, Drake LA, Adam AP, Schwartz BS, Singer HA, Swanson S, Mosher DF, Stewart R, Coon JJ, Jaitovich A. Large-Scale Multi-omic Analysis of COVID-19 Severity. Cell Syst. 2021 Jan 20;12(1):23–40.e7. doi: 10.1016/j.cels.2020.10.003. Epub 2020 Oct 8. PMID: 33096026; PMCID: PMC7543711.

45. Zhang Y, Zong L, Zheng Y, Zhang Y, Li N, Li Y, Jin Y, Chen L, Ouyang J, Bibi A, Huang Y, Xu Y. A single-cell atlas of the peripheral immune response in patients with influenza A virus infection. iScience. 2023 Nov 22;26(12):108507. doi: 10.1016/j.isci.2023.108507. PMID: 38089584; PMCID: PMC10711475.

46. Liao M, Liu Y, Yuan J, Wen Y, Xu G, Zhao J, Cheng L, Li J, Wang X, Wang F, Liu L, Amit I, Zhang S, Zhang Z. Single-cell landscape of bronchoalveolar immune cells in patients with COVID-19. Nat Med. 2020 Jun;26(6):842–844. doi: 10.1038/s41591-020-0901-9. Epub 2020 May 12. PMID: 32398875

47. Zhang Z, Zhang L, Wang K, Xie T, Zhang X, Yu W, Li Y, Shen L, Li R, Peng Z. Single-cell landscape of bronchoalveolar immune cells in patients with immune checkpoint inhibitor-related pneumonitis. NPJ Precis Oncol. 2024 Oct 5;8(1):226. doi: 10.1038/s41698-024-00715-6. PMID: 39369126; PMCID: PMC11455925.

48. Melms JC, Biermann J, Huang H, Wang Y, Nair A, Tagore S, Katsyv I, Rendeiro AF, Amin AD, Schapiro D, Frangieh CJ, Luoma AM, Filliol A, Fang Y, Ravichandran H, Clausi MG, Alba GA, Rogava M, Chen SW, Ho P, Montoro DT, Kornberg AE, Han AS, Bakhoum MF, Anandasabapathy N, Suárez-Fariñas M, Bakhoum SF, Bram Y, Borczuk A, Guo XV, Lefkowitch JH, Marboe C, Lagana SM, Del Portillo A, Tsai EJ, Zorn E, Markowitz GS, Schwabe RF, Schwartz RE, Elemento O, Saqi A, Hibshoosh H, Que J, Izar B. A molecular single-cell lung atlas of lethal COVID-19. Nature. 2021 Jul;595(7865):114-119. doi: 10.1038/s41586-021-03569-1. Epub 2021 Apr 29. Erratum in: Nature. 2021 Oct;598(7882):E2. doi: 10.1038/s41586-021-03921-5. PMID: 33915568; PMCID: PMC8814825.

49. Sun F, Guo ZS, Gregory AD, Shapiro SD, Xiao G, Qu Z. Dual but not single PD-1 or TIM-3 blockade enhances oncolytic virotherapy in refractory lung cancer. J Immunother Cancer. 2020 May;8(1):e000294. doi: 10.1136/jitc-2019-000294. PMID: 32461344; PMCID: PMC7254155.

50. Sun F, Li L, Xiao Y, Gregory AD, Shapiro SD, Xiao G, Qu Z. Alveolar macrophages inherently express programmed death-1 ligand 1 for optimal protective immunity and tolerance. J Immunol. 2021 Jul 1;207(1):110–114. doi: 10.4049/jimmunol.2100046. Epub 2021 Jun 16. PMID: 34135059; PMCID: PMC8674373.

51. Qu Z, Sun D, Young W. Lithium promotes neural precursor cell proliferation: evidence for the involvement of the non-conical GSK3β-NF-AT signaling. Cell Biosci. 2011; 1(1):18.

52. Qing G, Yan P, Xiao G. Hsp90 inhibition results in autophagy-mediated proteasome-independent degradation of IκB kinase (IKK). Cell Res. 2006; 16(11):895–901.

53. Qing G, Yan P, Qu Z, Liu H, Xiao G. Hsp90 regulates processing of NF-κB2 p100 involving protection of NIK from autophagy-mediated degradation. Cell Res. 2007; 17(6):520–530.

54. Yan P, Qing G, Qu Z, Wu CC, Rabson A, Xiao G. Targeting autophagic regulation of NF-κB activation in HTLV-I transformed cells by geldanamycin: Implications for therapeutic interventions. Autophagy. 2007; 3(6):600–603.

55. Xiao G. Autophagy and NF-κB: fight for fate. Cytokine Growth Factor Rev. 2007 Jun-Aug;18(3-4):233–43. doi: 10.1016/j.cytogfr.2007.04.006.

